# Gonadal remodelling in socially driven sex change is associated with novel and known sex genes, epigenetic reprogramming, and inflammation-mediated apoptosis

**DOI:** 10.64898/2026.09.03.749291

**Authors:** Chloé A. van der Burg, Alexander Goikoetxea, Simon Muncaster, Oscar Ortega-Recalde, Tim Hore, Kaj Kamstra, Olivier Fedrigo, Erich D. Jarvis, Alan Tracey, Kerstin Howe, Erica V. Todd, Neil J. Gemmell

## Abstract

Many fish undergo dramatic, socially cued female-to-male sex change, yet the events that initiate and then regulate gonadal reprogramming and reorganisation during this phenomenal metamorphosis are broadly unknown. The New Zealand spotted wrasse, *Notolabrus celidotus*, is a temperate fish species that displays the extraordinary ability to undergo protogynous sex change. Removal of a terminal-phase male from a social group triggers the most dominant female to change sex and become a male. To characterise the molecular changes associated with gonadal metamorphosis in spotty, we used genomic and transcriptomic approaches, generating a high-quality genome and a transcriptomic time-series that captures the sex change process from start to finish. These data, together with paired histological data, reveal distinct transcriptional profiles that characterise the process of sex change. As expected, the expression of masculinising genes steadily increases, while feminising genes steadily decrease, throughout the transition. We further identify novel candidate genes, including genes involved in immune signalling and tissue remodelling through inflammation-mediated apoptosis, whose expression strongly correlates with key events in sex change, while also confirming the roles of known sex determination and differentiation genes in this process. Collectively these data give us new insights into how a normally committed developmental process remains plastic and is reversed to completely alter organ structures, and also sheds light on the evolution of sex determination in other animals. This work furthers the development of the spotty as a new tractable model for sex change research.

## Introduction

The ability to change sex as an adult is a remarkable example of phenotypic plasticity. Over 450 species of fish can change sex, with the majority (∼66%) of these being protogynous sequential hermaphrodites (female-to-male) (Kuwamura et al. 2020). Well-known examples of fish that change sex include the protandrous (male-to-female) clownfish *Amphiprion spp.* (Avise and Mank 2009; Kuwamura et al. 2020), and commercially important protogynous species such as blue cod (*Parapercis colias*) and barramundi (*Lates calcarifer*) (Kuwamura et al. 2020). However, despite sex change being widespread among fishes, the molecular events that initiate and govern this phenomenon are only just now beginning to be investigated (Zhou and Gui 2010; Todd et al. 2019; Ortega-Recalde et al. 2020; Goikoetxea et al. 2021; Muncaster et al. 2023; Quertermous et al. 2025).

The New Zealand spotty wrasse (paketi, *Notolabrus celidotus*) is an emerging model for protogynous sex change (Goikoetxea et al. 2020; Goikoetxea et al. 2021; Kamstra et al. 2023; Muncaster et al. 2023; Quertermous et al. 2025). It is a temperate species, endemic to New Zealand, and is found ubiquitously and abundantly along the coastline. Spotty wrasse have two dimorphic and diandric male phenotypes, known as initial phase (IP) males and terminal phase (TP) males (Jones 1980; Robertson 2020). The majority of juveniles sexually mature as IP females, with a small percentage (5-10% in the wild) differentiating as IP males (Jones 1980), once they reach approximately 100-110 mm standard length (Hamer et al. 2025). IP females and IP males are virtually externally indistinguishable from each other, but TP males are easily distinguished from IPs by differences in colouration and markings (Fig. S1). Sex change in spotty wrasse is socially-controlled; the most dominant (and usually largest) female will change sex to male upon removal of the TP male (Goikoetxea et al. 2021; Quertermous et al. 2025). Spotty wrasse are amenable to captivity and sex change is easily induced in the lab; the transition takes approximately 60 days to complete. Sex change has been observed and can be induced throughout the year (Thomas et al. 2019; Goikoetxea et al. 2021), although sex change occurs more readily and rapidly after breeding season (around Austral spring, in August-November), and occurs around November to May in the wild (Jones 1980).

Sex determination mechanisms are diverse across taxa, however, the suite of master sex-determining genes employed tend to be conserved, with conserved physiological roles (Herpin and Schartl 2011; Cutting et al. 2013; Capel 2017; Bertho et al. 2021; Kitano et al. 2024). These ‘classic’ genes include well-documented masculinising genes such as *sox9*, *amh*, *cyp11c1*, *dmrt1* and *hsd11b2*, and feminising genes such as *foxl2*, *cyp19a1a*, *rspo1*, *wnt4* and *ctnnb1*. Some of these genes have a direct effect on physiological sex via the synthesis of estrogens or androgens, for example, *cyp19a1a/b* or ‘aromatase’ converts androgens into estrogens and *cyp11c1* catalyses cortisol to 11-ketotestosterone (11-KT). Many of these genes are therefore also key investigatory targets in studies exploring adult sex change in teleosts, *e.g.,* in the bluehead wrasse (*Thalassoma bifasciatum*) and in previous work on spotty wrasse (Thomas et al. 2019; Todd et al. 2019; Ortega-Recalde et al. 2020; Goikoetxea et al. 2021; Muncaster et al. 2023).

Epigenetic regulation is an important part of sex determinism and sex plasticity in fish and other vertebrates (*e.g.,* bearded dragon) (Shao et al. 2014; Deveson et al. 2017; Piferrer 2018; Ortega-Recalde et al. 2020; Piferrer 2021). In bluehead wrasse, global DNA methylation (DNAm) increases in the gonads as maleness increases (Todd et al. 2019), which is similarly reflected in spotty wrasse through the increase in both epigenetic regulatory factor expression (Muncaster et al. 2023) and global DNAm in males vs females (Robertson 2020), although notably, DNAm has not yet been directly measured in spotty wrasse transitional gonads. Interestingly, DNAm is known to be influenced by environmental factors (temperature, salinity, season) and in spotty wrasse, global DNAm levels are higher in female ovaries during spawning season, than outside of spawning season (Robertson 2020). Other species, such as the protandrous Barramundi (*Lates calcarifer*), also exhibit environmental differences in DNAm. One study (Budd et al. 2022) found region-specific (coastline) differences in the DNAm of four sex-determining genes in barramundi, as well as expected sex-specific DNAm, *i.e.,* male genes are hypermethylated in females and vice versa, which drives the suppression of the opposite sex phenotype.

Currently there are few full time-course transcriptomic studies of sex change in protogynous fish gonads, and investigation of multiple transitional states is particularly lacking. One notable exception is work on the bluehead wrasse (Todd et al. 2019) which investigated seven transitional stages, as well as female and male samples. Other studies investigating protogynous (female-to-male) gonadal transition include ricefield eel *Monopterus albus* (Fan et al. 2022; He et al. 2022) and a meta-analysis of seven teleost species (Nozu et al. 2024). Previous work focussing on specific genes using qPCR (Thomas et al. 2019) or probe-based detection of expression (*i.e.,* nanoString nCounter^TM^) (Goikoetxea et al. 2021; Goikoetxea et al. 2022; Muncaster et al. 2023) has been undertaken in the spotty wrasse.

Here, we present a chromosome level genome for the spotty wrasse, along with the first full transcriptomic analysis of gonadal sex change. We investigate, using multiple bioinformatic approaches, the full suite of genes expressed throughout each stage of sex change, from female, through multiple transitional stages to terminal phase male (TP male), as well as initial phase (IP) males, which can undergo role change to TP males. These data give us novel insight into potential early and consistent drivers of sex change in spotty, as well as unique patterns of gene expression that typify each gonadal phenotype.

## Results

### A high-quality spotty wrasse reference genome

We assembled the genome of a male individual, following the Vertebrate Genomes Project pipeline v1.6 (Rhie et al. 2021). The ID of the primary chromosomal level assembly is fNotCel1.pri (GenBank accession GCA_009762535.1) and the alternate contig-only assembly fNotCel1.alt. The total primary assembled genome size was 846.7 Gb, within the range we find for teleost fish species. Our genome shows solid assembly metrics, with the contig N50 at 3.7 Mb and the scaffold N50 at 37.1 Mb, above the minimum standards of 1 Mb and 10 Mb set for a high-quality, vertebrate, reference genomes (Rhie et al. 2021), respectively. We curated and identified 24 chromosomes, as we have seen in other teleost fish species, and almost all sequence (97.6%) mapped to chromosomes. Our genome assembly registered a BUSCO score of 96.2% complete genes. We also sequenced and assembled the complete mitochondrial genome (accession CM019797.1), which was 16,525 bp in size.

### Gonad-wide gene expression changes during sex change

We performed RNA-sequencing on a time series sampling of spotty wrasse gonads following (chemically or socially) induced sex-change, utilising 96 samples from across four experimental sampling batches. While each of the four experimental sampling batches across 2014, 2015, 2016, and 2018 were designed with different parameters concerning season, sampling methodology and sex-change induction method (see methods section ‘Experimental design and sample collection’), together these experiments captured gonadal samples of the entire sex spectrum in spotty wrasse from female, through three transitional states (early, mid, late), to terminal-phase male. Multiple other sex phenotypes and states were also captured, including IP males, breeding and non-breeding females. These data are investigated herein to determine how gene expression is driving sex change in the gonad and what factors most strongly are associated with each gonadal sex type.

### Dataset parameters, read and count processing

Reads are available at NCBI BioProject PRJNA631152. After trimming, the average number of reads was 11,013,818 (range of 6,804,419 - 16,267,816) per library and the overall quality score for all reads is >Q30 (Fig S3). An average of 86.15% of reads uniquely mapped to the genome (range of 72.81 % -91.06 %), see Fig. S4a. Most libraries showed good 5’->3’ gene coverage, see Fig. S4b. STAR detected 28,261 gene IDs (all genes, no pseudogenes) and after filtering to remove very low count genes (removing genes with counts less than 10 in most samples), 22,231 genes remained. Counts were then further ‘group-based’ filtered (see explanation in Supplementary Materials 3d. Group Filtering), to retain a 6-trait group (F, ET, MT, LT, TPM, IPM; female, early-transitioning, mid-transitioning, late-transitioning, terminal-phase male, initial-phase male) matrix resulting in 18,320 genes, with lowly expressed genes across all trait groups removed. Sample metadata and gene expression profiles for each trait group were explored and visualised in Figure 1. These data show number of samples in each trait group by experimental batch, with the greatest number of samples obtained in the ET group (n=28), and the least in IPM (n=4) (Fig.1a). Investigation of sample variation in lower-dimensional space shows that sex-change is a continuous spectrum from female to male (Fig. 1b).

**Figure 1.**
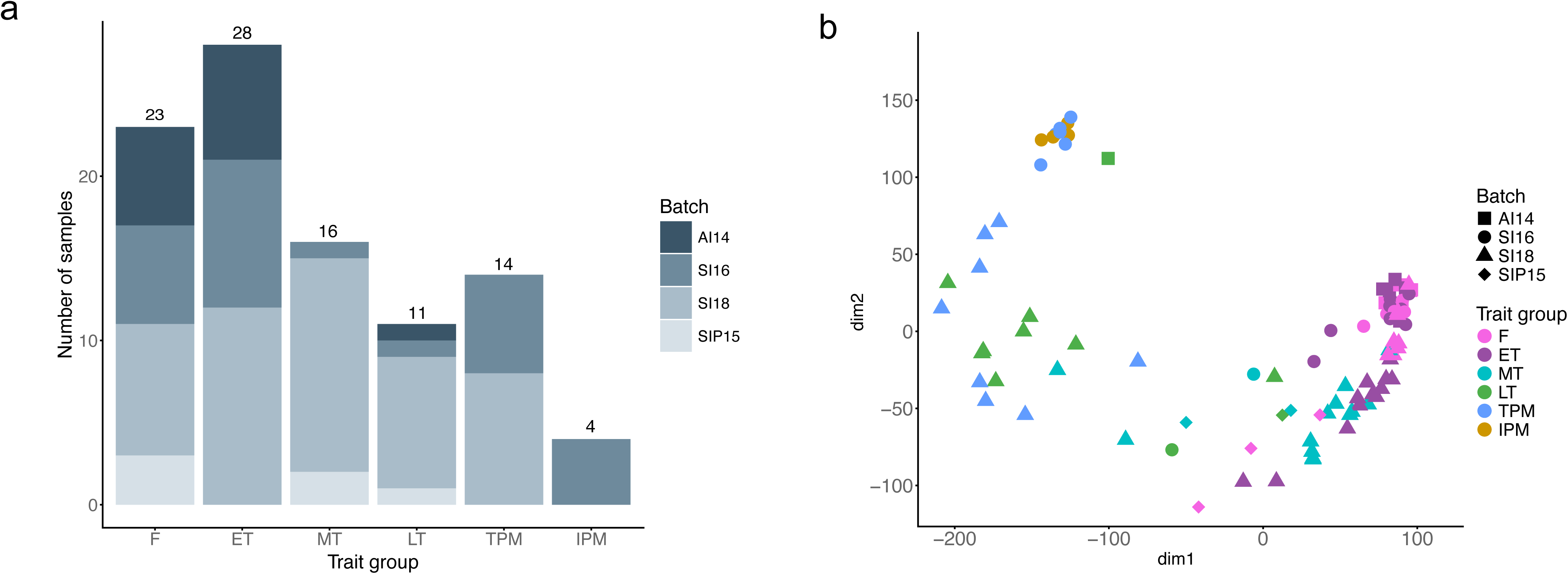
Exploration and visualisation of dataset. A) Distribution of sample histology designations (*i.e.,* trait groups) across batches. B) GLM PCA (“group-filtered”, variance stabilised DESeq2 counts). Batch refers to four separate experiments, see Supplementary Material for further explanation. F = Female, ET = Early-transitioning, MT = Mid-transitioning, LT = Late-transitioning, TPM = Terminal-phase male, IPM = Initial -phase male.

### Differential gene expression analysis finds progressive activation of male-biased genes

Differential gene expression (DGE) analysis was performed using 6 trait-groups (F, ET, MT, LT, TPM, IPM), comparing all trait groups against females. In total, 43,737 instances of DE were detected (across all up-regulated and down-regulated genes vs F), however of these, 15,772 unique genes were DE, indicating most of the genes in the dataset were DE. Differential expression increases as sex change proceeds from female to male, with the least number of DE genes detected when comparing F vs ET (64 DEGs), and the highest number of DE genes detected when comparing F vs TPM (13,730 DEGs) (Table S2a). The number of genes uniquely upregulated also increases through sex change (or towards maleness) (Fig. 2a and Fig. 2b), although this pattern does not hold for downregulation, with MT samples showing the most (n=757) uniquely downregulated genes compared to females (Fig. 2c). We detected 36 genes that are always upregulated in all traits compared to females (Fig 2a-b, Supplementary Fig. S12), suggesting a potential role in initiating and maintaining sex change towards a male state. These 36 genes have roles in several major processes, including apoptosis, neuroendocrine and signalling functions, structural/ECM, immune responses and epigenetics. No genes were found to be consistently downregulated (Fig. 2c). Expression patterns of all upregulated genes were further explored (Fig. 2 d.i-h.i), showing that as gonads transition through from female to male, gene function changes from largely processes involved in transcription, RNA processing and female reproductive processes, through to an increase in genes related to immune processes, neural activity, tissue patterning and spermatogonia proliferation (Fig 2d.ii-h.ii; Supplementary Material section ‘DGE’; Supplementary Tables S3-S8). Both states show expected biases in respective reproductive terms (*i.e.,* egg formation in females, sperm proliferation in males), confirming dataset integrity. We find a significant increase in the GO term “immune response” during the transition to male, beginning at the mid-transitional stage (*e.g.,* Fig 2e-g.ii), highlighting a potential novel role of the immune system in male/male-biased gonads. We also find enriched GO terms “extracellular space” and “membrane” appearing consistently as top GO terms for MT, LT and TPM in DGE analysis (Fig. 2e-g.ii), further highlighting the massive structural changes the gonadal tissue undergoes as it transitions from female to male.

**Figure 2.**
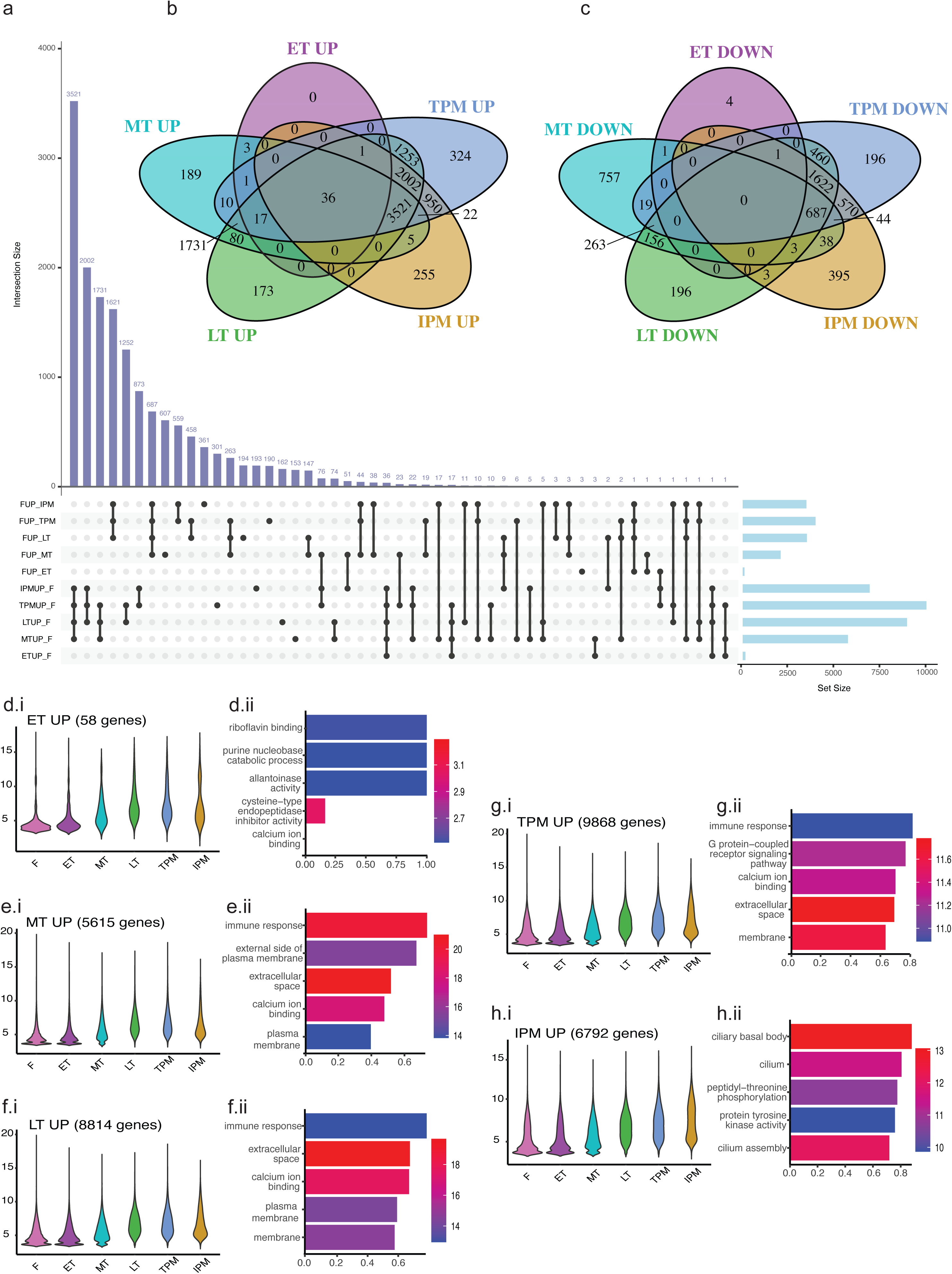
Differential gene expression comparing females to all traits. a) Venn upset diagram showing all genes upregulated and downregulated in each trait vs female. Pairwise comparisons are indicated on the left as “Trait Up/down, vs trait”: *e.g.,* FUP_ET = All genes upregulated in F (female), when compared to ET (early-transitioning). Intersections with 0 genes up/downregulated not shown, all gene intersections detailed in Supplementary Table S2b; b) Venn pie showing all genes upregulated in each trait, vs female, including 36 genes (centre of Venn) that are DE in all traits compared to females; c) Venn pie showing all genes downregulated in each trait, vs female; d-h.i) Expression of genes upregulated in ‘X’ trait group, across all traits; d-h.ii) Top 5 enriched GO terms for the gene set (lowest p-value). For d-h, gene expression is shown as variance stabilised DEseq2 counts. Colour legend for top 5 GO terms is shown as -log(10) pvalue and x-axis shows the Gene Ratio (numDEInCat/numInCat).

### Distinct gene expression profiles typify each sex phenotype and transitional state

After group-based filtering (6 trait-groups) the DESeqDataset to remove low count genes, 18,320 genes remained in total and were used for further weighted gene co-expression network analysis (WGCNA), to find genes that correlate with each distinct sex phenotype or transitional state. WGCNA outlier gene detection was initially run and found 1,682 outlier genes (*e.g.,* missing entries, zero-variance genes), which were removed, but further group-based filtering retained the same genes (*i.e.,* 18,320) as when no outlier detection was performed, indicating it was redundant. A soft power of 20 was chosen based on visualization of the scale free topology and mean connectivity (Fig. S13).

Nine module eigengenes (MEs) were detected after merging (Supplementary Fig. 14a-b), with the smallest containing 245 genes (magenta) and the largest containing 4944 genes (turquoise); 3909 genes were unassigned (*i.e.,* “grey” module) (Supplementary Tables S9a-b). Correlation analysis shows how correlated each module is with each trait group, with a highest positive correlation of 0.61 (MEblue with TPM) and a lowest negative correlation of -0.58 (MEbrown with TPM) (Fig.14c). The expression of genes within modules across traits was also explored (Fig.3d), and generally, changes in gene expression across traits appears to be subtle, however, in all modules there is a significant statistical difference between F gene expression and TPM gene expression (Supplementary Table S9c). Correlation analysis also shows there are multiple distinct patterns in gene expression across the transition (Fig. 14c). Genes in modules ‘pink’ and ‘red’ show strong mid-transitional changes. Genes in the ‘green’ module are most highly positively correlated with IPM. Masculinising trends can be seen clearly in modules ‘turquoise’ and ‘blue’ (Fig.3c; correlation with TPM=0.5 and 0.61, respectively), and feminising trends can be seen clearly in modules ‘yellow’ and ‘magenta’ (Fig.14c; correlation with female=0.48 and 0.5, respectively). Detailed description of these modules are in Supplementary Materials (<u>6. WGCNA</u> section).

To further hone in on which genes are driving each trait group during sex change, the expression of the top 30 positively and top 30 negatively correlated genes (full list of gene-trait correlations in Supplementary Table S11) were plotted as boxplots in Figure 3, alongside the top 5 GO annotations (full GOseq results in Supplementary Table S12). Distinct gene expression patterns can be observed (Fig. 3a-f.i; 3a-f.iii) and together with gene ontology results (Fig. 3a-f.ii; 3a-f.iv), indicate that each trait group has a unique signature of gene expression that typifies the trait. Gene sets are all uniquely correlated with each trait (*i.e.,* there are no repeated genes across sets for each correlation and trait combination) and generally gene expression correlates higher with males (Fig. S16).

**Figure 3.**
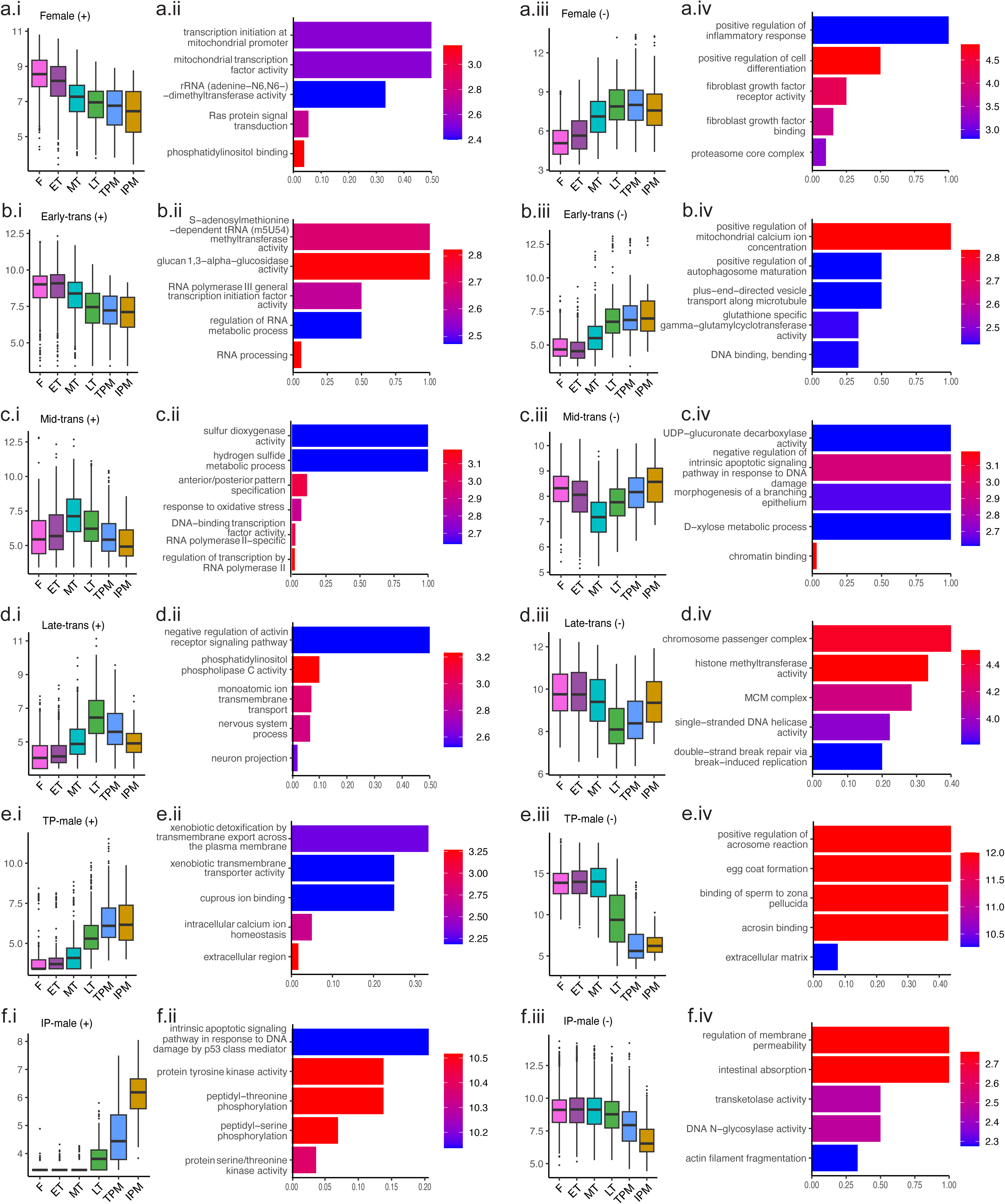
Gene-trait correlations. The expression of the top 30 genes most highly correlated with each trait are plotted, showing the change in expression across all traits. For each set of 30 genes (positively and negatively correlated), the top 5 enriched GO terms are shown, with the gene ratio plotted on the x-axis and the colour indicating the -log_10_Pvalue (*i.e.,* red = lower P-value, blue=higher p-value). There is no overlap between gene sets; each set of 30 are unique. Expression is shown as variance stabilised DESeq2 counts. (i): top 30 genes positively correlated (+) with the indicated trait and the associated top 5 enriched GO terms in (ii). (iii) top 30 genes negatively correlated (-) with the indicated trait and the associated top 5 enriched GO terms in (iv).

### Investigation of classic and putative sex change genes show cohesion with the literature

Classic and putative sex change genes, identified from key literature (references in Table 1 in Materials and Methods), were plotted in spotty to investigate change in expression over trait groups (Figure 4; expression values in Supplementary Table S13.). Statistical analysis comparing ‘Female’ versus ‘TP male’ samples show significant differences in gene expression for most genes (Supplementary Table S14 shows ANOVA results). Twelve genes show a clear female bias (p-value ≤ 0.001): *actb1* (HK), *foxl2a, ctnnb1*, *znrf3*, *dnmt1*, *h2az1*, *pou5f3*, *bmp15*, *tp53*, *cyp19a1a*, *gdf9* and *figl1*. One gene, *wnt4*, shows a slight (p-value ≤ 0.01) female bias. Twenty-two genes show a clear male bias (p-value ≤ 0.001): *g6pd* (HK), *dmrt1*, *sox9a*, *sox8a, gata4*, *wnt4b*, *rspo1*, *fancl*, *dnmt3bb.1*, *dnmt3ba*, *dnmt3ab*, *tet1*, *tet3*, *suz12b*, *csf1ra*, *csf1b*, *irf8*, *cyp11c1*, *hsd11b2*, *nr3c2*, *gsdf* and *amh.* Six genes show a slight (p-value ≤ 0.01) male bias: *kdm6bb*, *tet2*, *eed*, *ezh2*, *il34* and *nr3c1*. There is no significant difference between female and TP male samples for three genes: *jarid2b*, *chek2* and *nr0b1*. *Jarid2b* shows lower expression in MT samples, which is also consistent with bluehead wrasse (Table 1), which shows mid-sex change suppression of this gene (statistical difference detected between MT vs F and MT vs TPM; p-value for both comparisons ≤ 0.001). While generally the expression of epigenetic regulators trends up in males, we find the genes *ezh2* and *jarid2b* to be suppressed at mid-transition, with expression significantly increasing again in TP males for *ez2h,* but not for *jarid2b* (see Table S14, Fig. 4d-e), a finding supported by work on bluehead wrasse (Todd et al. 2019) and earlier work on spotty (Muncaster et al. 2023).

**Figure 4.**
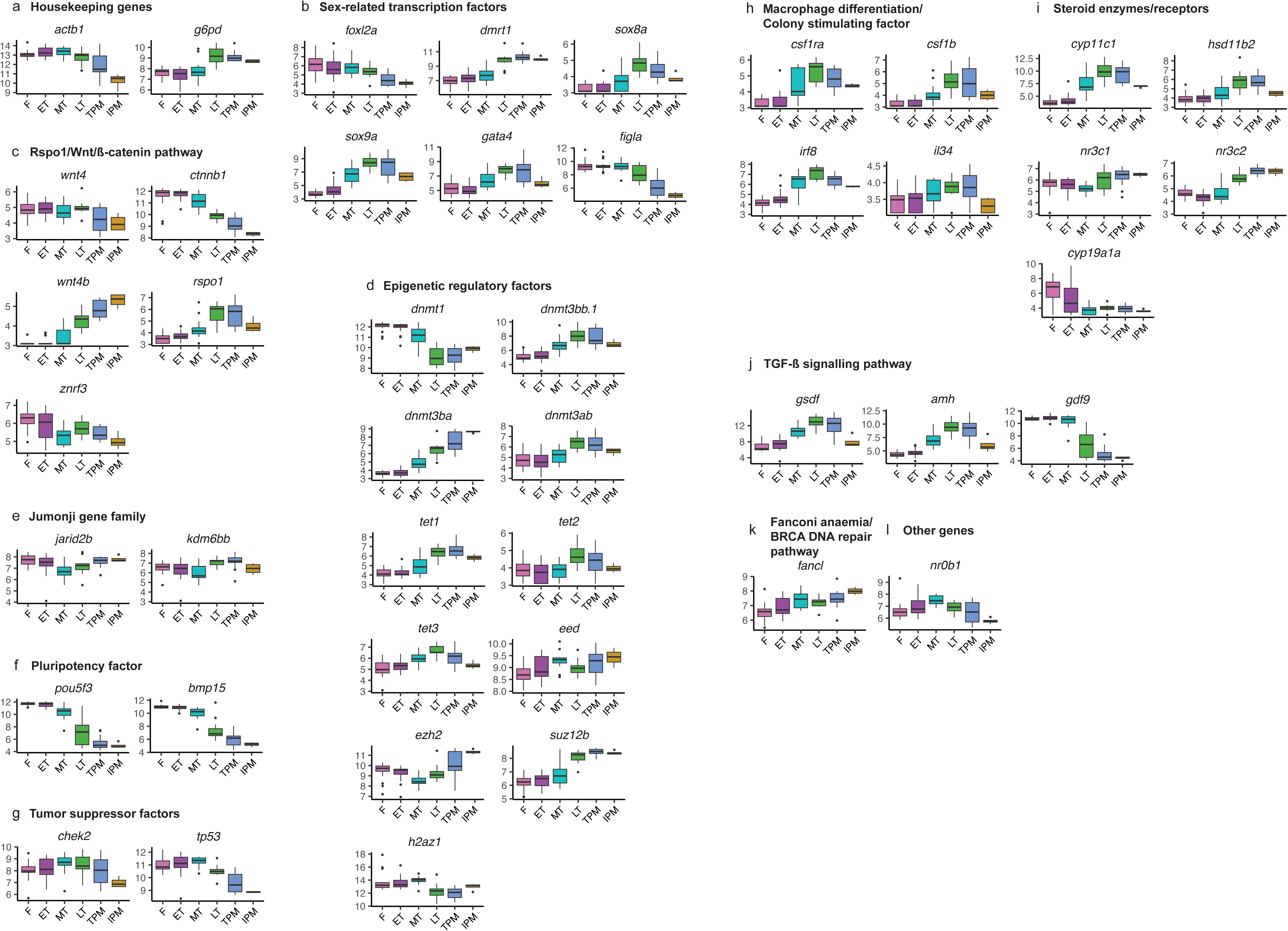
**Expression of classic and putative sex change genes across trait groups**. Expression on the y-axis is shown as variance stabilised DESeq2 counts. Trait groups are shown left to right on the x-axis as female transitioning through to male, as follows: F= female; ET = Early-transitioning; MT = Mid-transitioning; LT = Late-transitioning; TPM = Terminal-phase male; IPM = Initial-phase male. Refer to Table 1 for gene descriptions.

**Table 1.**
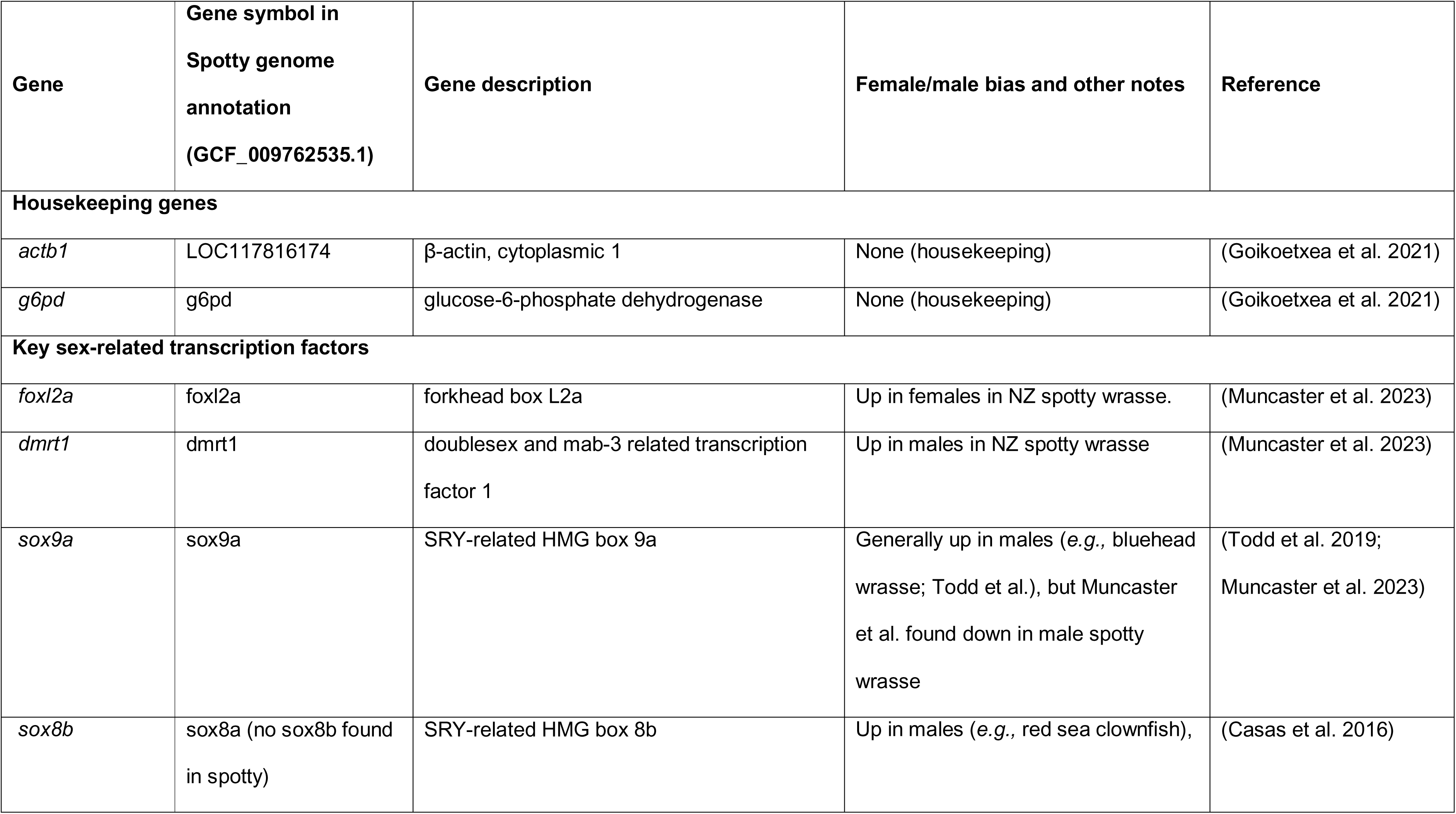
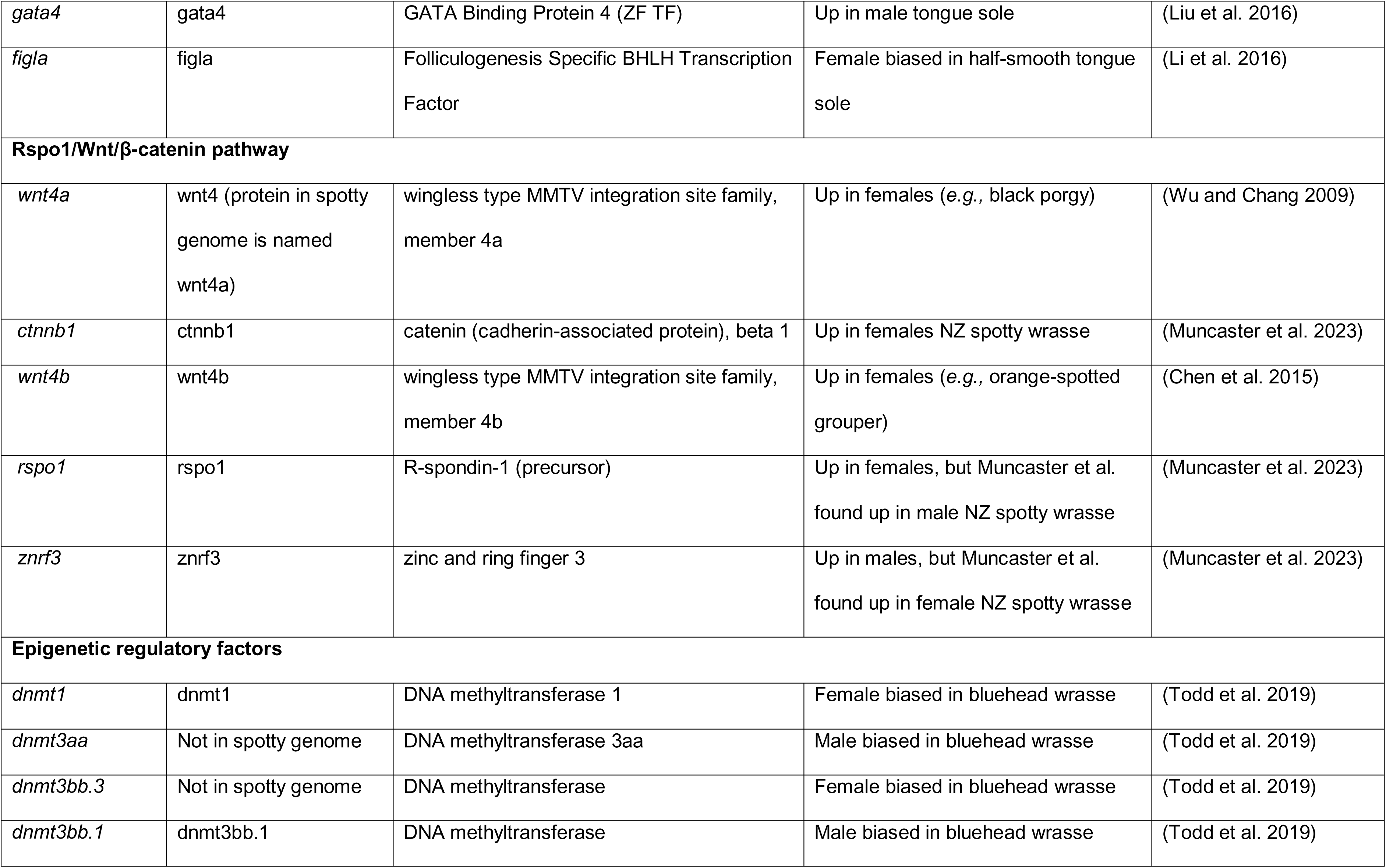
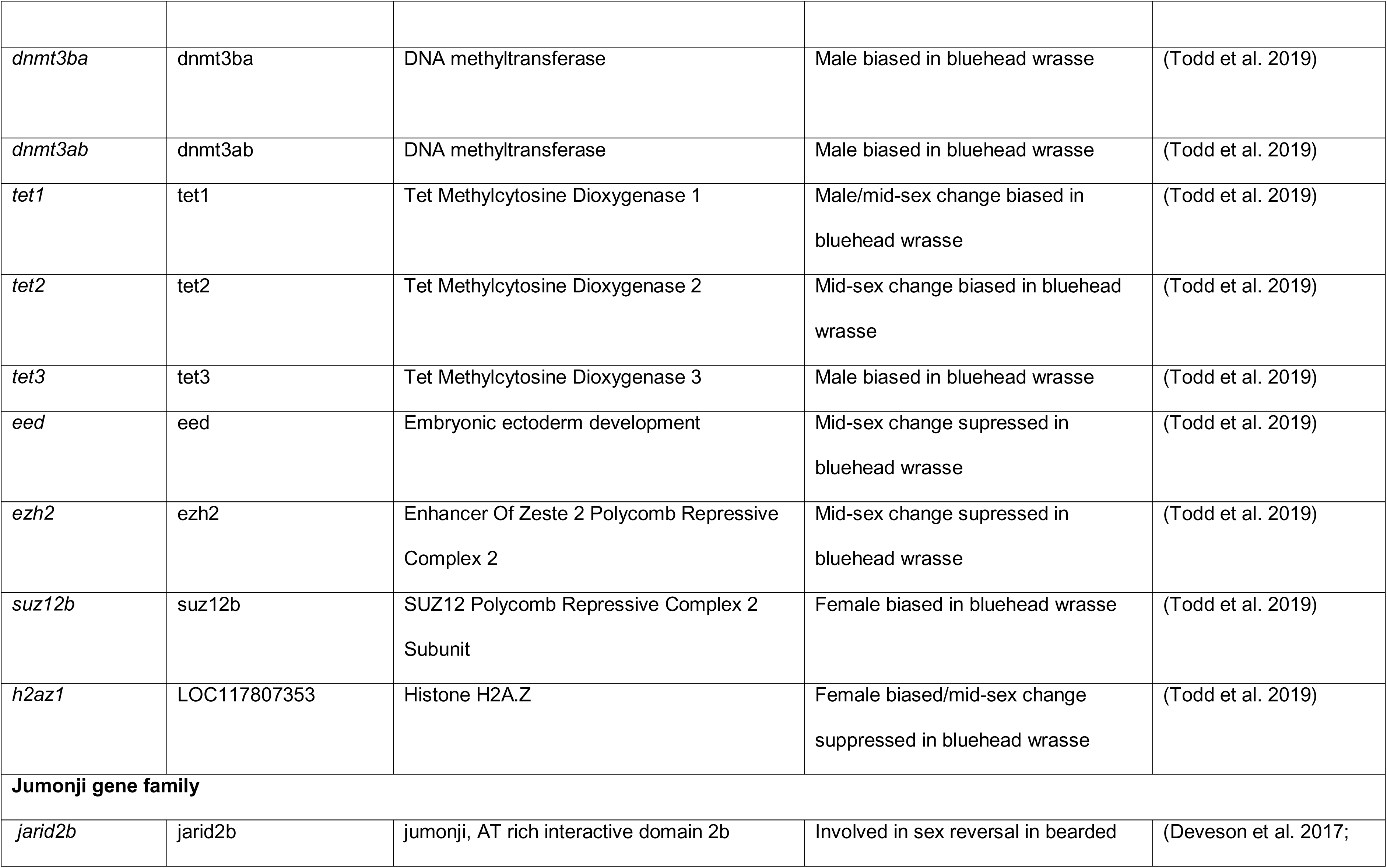
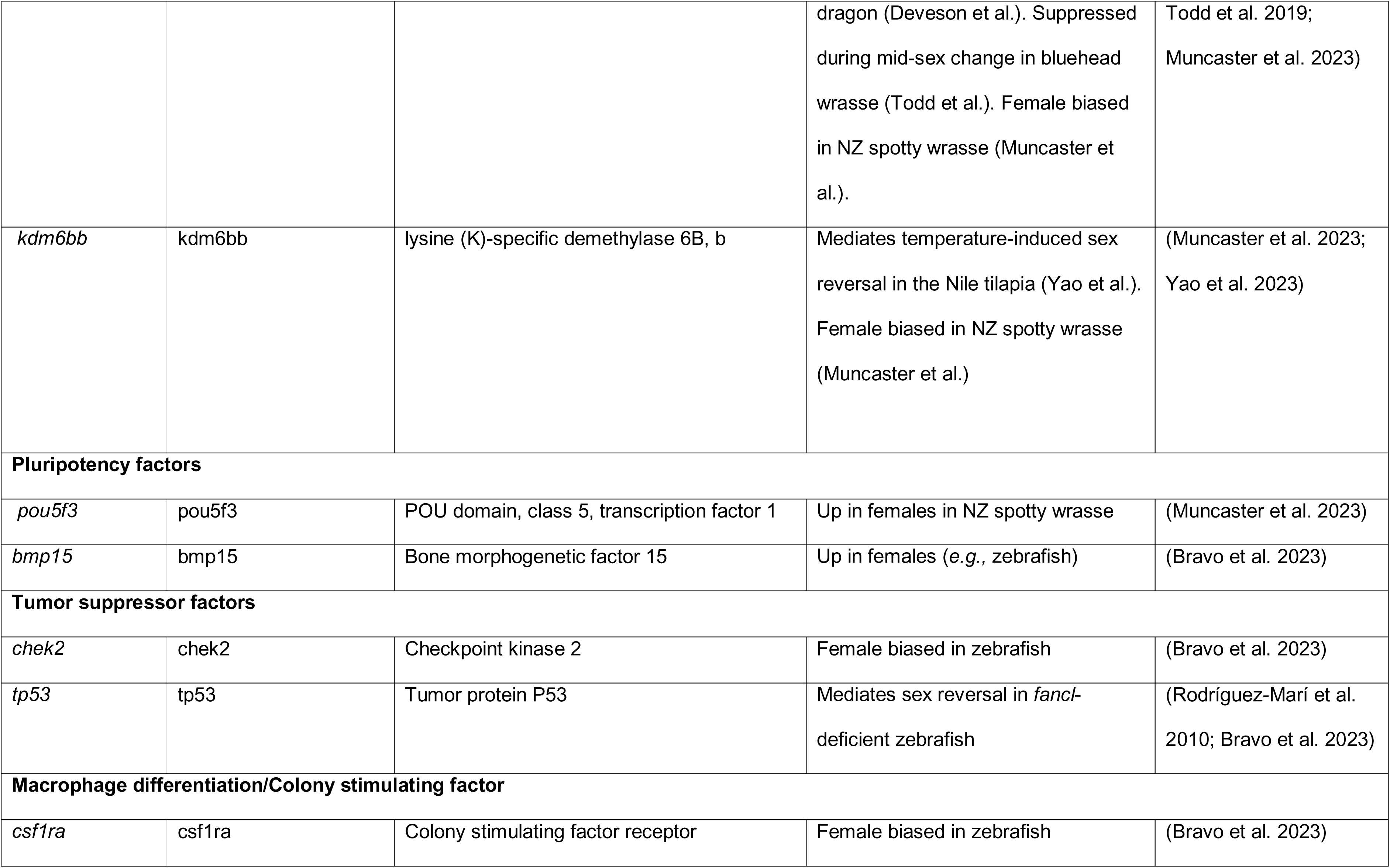
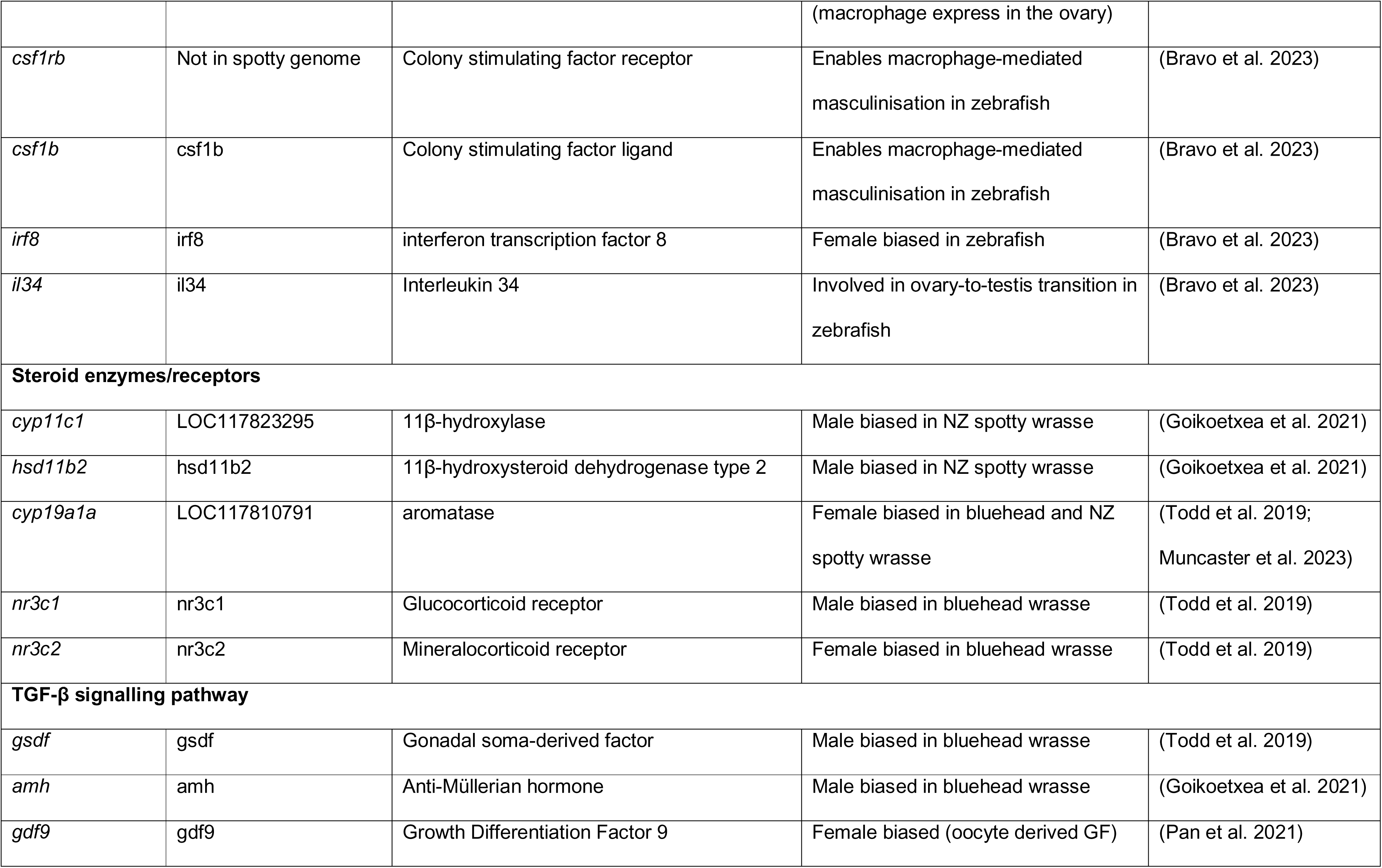
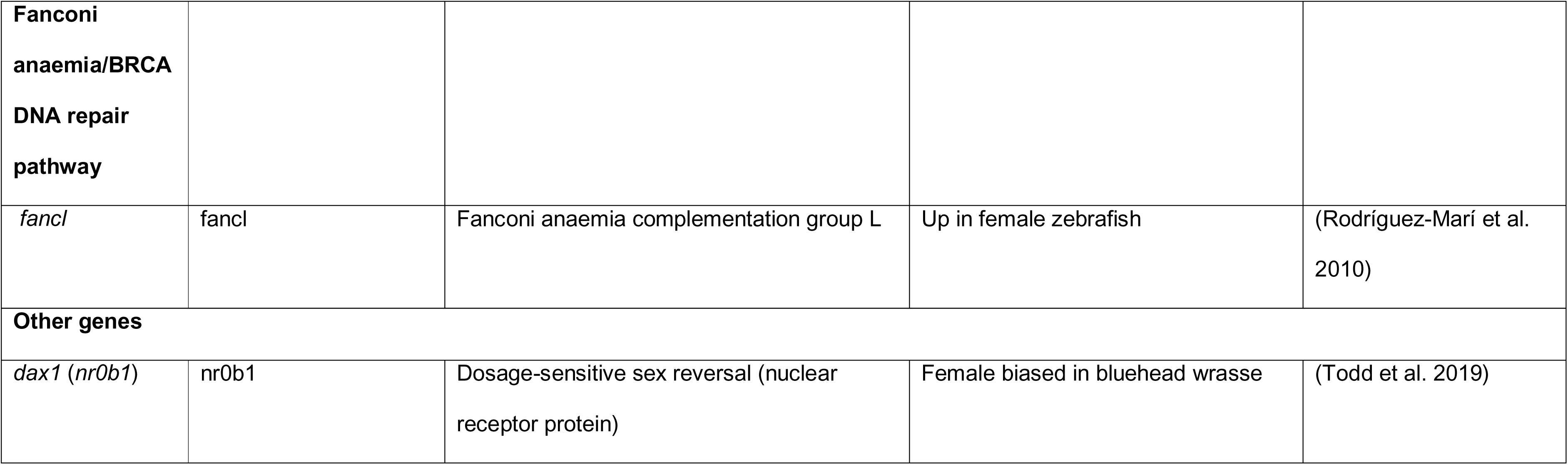
List of classic and putative candidate sex-change genes. Notes on female/male bias are from references cited and does not necessarily reflect results found here for spotty.

## Discussion

The process of socially-controlled sex change represents a striking example of phenotypic plasticity in response to environmental and social cues. Herein we show that distinct gonadal states during sex change in the New Zealand spotty wrasse are characterised by unique transcriptional profiles. Many classic sex-determination and differentiation genes show expression patterns consistent with previous studies, while numerous genes not previously highlighted in sex-change research are strongly associated with particular gonadal states. We also identify 36 genes that are consistently upregulated across non-female gonadal phenotypes and therefore represent candidate components of a broader masculinisation-associated transcriptional programme.

### Each phenotypic state is typified by a unique gene suite and physiology

Each of the sex phenotypes (female, initial-phase male, terminal-phase male) and transitional state phenotypes (early, mid, late) described here in spotty show a unique pattern of gene expression that exemplifies different cellular and molecular characteristics, corresponding to a unique physiological state. Broadly, females are associated with increased transcription and RNA processing, and lipid/cholesterol biosynthesis and processing. On the other end of the spectrum, TP males exhibit gene expression associated with increased cell signalling, membrane protein trafficking, immune response, and neural and synaptic signalling. The point where gene expression switches from a female-biased profile to a male-biased profile appears to be between mid and late transition, with most MT samples having a female-biased profile, and LT samples a more male-biased profile. This switch accompanies a change in cellular and molecular function from a focus on internal cellular mechanics to overhauls in tissue patterning and structure. Both female and TP males show consistent profiles with the protogynous bluehead wrasse (Todd et al. 2018; Todd et al. 2019), indicating general patterns of conserved sex change physiology, despite large differences in time line for sex-change between the two species *i.e.,* the bluehead wrasse completes sex change in 8-10 days; spotty requires approximately 60 days.

Steroidogenic factors are generally key maintainers of binary sex state once achieved in both protandrous (Casas et al. 2016; Wang et al. 2022) and protogynous fish (Nozu et al. 2024), regardless of direction of sex change, in line with gonochoristic sex-specific steroid biosynthesis and maintenance. Interestingly, while both IP and TP males have similar functioning gonads (*i.e.,* sperm-producing), which may bring the expectation of similar transcriptional profiles, we find that IP males and TP males gonads are transcriptionally both unique male phenotypes, in line with literature on ‘sneaker’ (subordinate) vs ‘bourgeois’ (dominant) bluehead wrasse males (Todd et al. 2018). IP males are associated with increased phosphorylation, kinase activity, and apoptosis, as well as many sperm-associated terms (same as TP males), but do not show the increased immune response term that is seen TP males. These findings further exemplify the uniqueness of each of the three sex phenotypes in spotty, alongside other research showing that gross gonadal morphology is also very different, *i.e.,* IPs have solid testes, TPs have hollow testes (Robertson 2020). Future work could employ the genes identified here as a potential diagnostic tool for confirming the state of the gonad. Further work should continue to refine the stages or timepoints along the sex change spectrum, especially focussing on refining more subtle markers or changes in the mid-transition stage.

### Genetic drivers of gonadal sex change

Increase in transcript diversity

Sex change in spotty gonads is driven by an increase in transcription as females transition to male, with a steady increase in both the number of expressed genes (*i.e.,* transcript diversity) and the number of differentially expressed genes in each trait. As expected, the largest number of differentially expressed genes was between females and TP males. This is highlighted by the differential expression of transcription factors such as *sox9a*, *etv1*, *foxc1a*, and *nkx3.3*, which are central to the reprogramming of gonadal fate and are consistently upregulated in all trait groups compared to females. *Sox9* in particular is a well-established driver of testicular differentiation in vertebrates, and in protogynous species (such as many wrasses, including spotty), *sox9* expression coincides with the initiation of spermatogenesis and suppression of ovarian function (Herpin and Schartl 2011; Ortega-Recalde et al. 2020; Vining et al. 2021).

Early and consistent genetic drivers of sex change

Apoptosis and cell remodelling appear are key processes in sex change, emphasising the dual role of cell death and survival during sex change. Apoptosis is critical for clearing ovarian follicles and creating space for the formation of testicular tissue and previous studies in other protogynous fish have shown that apoptosis of oocytes and granulosa cells is a hallmark of ovarian regression (Morais et al. 2012; Liu et al. 2017; Fan et al. 2022).

The upregulation of neuroendocrine and signalling genes, highlights the significance of endocrine and intracellular signalling in mediating socially-controlled sex change. GABA receptors mediate inhibitory signalling in the nervous system and are known to influence the hypothalamus-pituitary-gonadal (HPG) axis, which controls gonadal sex steroid production (Kamstra et al. 2024). The increase in calcium transporting and binding genes such as *calml4a* and *PMCA*-like indicate that intracellular Ca^2+^ signalling is orchestrating the change (Strehler et al. 2007), with some work linking calmodulin to the regulation of androgen production in teleosts (He et al. 2022). Structural and ECM related genes also appear frequently during sex change, which is as expected, given the massive amount of tissue remodelling required to convert a gonad from female to male.

Interestingly, we observed enrichment of immune and inflammatory processes beginning during the mid-transitional stage, a pattern that is not frequently reported in other fish sex-change studies (Todd et al. 2019; Ortega-Recalde et al. 2020; Muncaster et al. 2023; Nozu et al. 2024). The relative absence of this bias in IP males is consistent with a transition-associated rather than simply male-associated signature. These data identify immune signalling and apoptosis as candidate contributors to gonadal remodelling, but functional studies will be required to determine whether these processes have a causal role in sex change.

Lastly, epigenetic regulation has emerged as a common mechanism of sex determination and differentiation across taxa with plastic sex systems (Capel 2017; Piferrer et al. 2019; Ortega-Recalde et al. 2020; Piferrer 2021). Here, we observed substantial changes in the expression of genes involved in DNA methylation, histone modification and chromatin regulation as gonads transitioned from female to male. These patterns are consistent with altered epigenetic regulatory activity during sex change and resemble patterns reported previously in bluehead wrasse (Todd et al. 2019). However, DNA methylation was not measured directly in transitional spotty gonads, and the direction or genomic targets of methylation changes therefore cannot be inferred from these transcriptomic data. While in spotty wrasse the methylation levels in gonads has not been specifically measured (Muncaster et al. 2023), the increase found herein in DNA methyltransferase genes in males, as well as other genes involved in histone modification and chromatin binding (*e.g., suz12b*, *ezh2*, *eed*, *tet1, tet2, tet3*), indicates that spotty likely are similar to bluehead wrasse in this regard. Interestingly, while methylation silences gene expression via suppressing transcriptome factor binding at promoter regions, we find more gene diversity and differential expression in males, indicating methylation here is likely playing a different role in male/male-biased gonads. Beyond methylation, other epigenetic changes such as histone deacetylation and subsequent chromatin remodelling are also implicated in sex-change (Ortega-Recalde et al. 2020) and herein we find many genes involved in all aspects of epigenetic modifications showing variable expression (Fig 5). Together, these changes suggest that sex change is underpinned by dynamic epigenetic reprogramming, allowing for a rapid and stable shift in gonadal identity. Here we investigated expression at a gene level, but studies show that much more fine-scale changes can catalyse the switch from female to male *e.g.,* an alternatively-spliced transcript variant of the gene *Kdm6bb* (histone demethylase) activates other male-biased genes downstream in Nile tilapia (Yao et al. 2023). For future work, we suggest research to focus on a few of the epigenetic targets identified here and investigate transcript or isoform dynamics using a more fine-scale analysis.

### Classic genes, novel genes, and the evolution of sex change

The evolution of plastic sex change systems is complex, and varied mechanisms have evolved throughout taxa, with no single conserved way to initiate the determination of sex (Capel 2017). In spotty, here we show many ‘classic’ genes are expressed in a conserved way, in line with the literature, and cases where expression differs to vertebrates, expression tends to be consistent with the more closely related bluehead wrasse. However, we find many genes that are not in this ‘classic’ list that are highly positively or negatively correlated with each stage. Of all the top 30 most highly positively and negatively correlated genes in each trait group (Fig. 3, n=360 genes), only one gene was also found in the ‘classic’ list (*nr3c2* - negatively correlated with ET). Further, of all the genes that are significantly upregulated in all stages against female (Fig 2b), 35 of these 36 genes are not found in the ‘classic’ list either (*sox9a* being the exception). Given the diverse nature of hermaphroditism in fish and the complex, interspersed evolutionary context (see Fig 1 of (Kuwamura et al. 2020) for phylogeny), coupled with the relative paucity of genomic resources exploring sex change in fish, it is difficult to confidently identify what is genuinely novel gene expression at this stage. Further comparative evolutionary and functional genomic analyses of these loci will be necessary to determine whether the observed expression dynamics in spotty represent lineage-specific innovations or reflect under-investigated, conserved mechanisms of sex reversal. While it is possible spotty achieve sex change using truly novel genes, it is likely that they re-deploy classic sex determination/differentiation pathways using a suite of regulatory mechanisms which may themselves be novel, and evidence of regulation of sex differentiation or sex reversal by miRNA, ncRNAs or alternative splicing in fish is beginning to emerge (Lu et al. 2022; Tang et al. 2022; Zhang et al. 2022; Jiang et al. 2024; Geffroy et al. 2025; Lin et al. 2025).

Finally, seasonality is likely to impose a strong background transcriptional structure on the gonads of the temperate spotty wrasse that is largely absent in the tropical bluehead wrasse. In spotty wrasse, gonadal state is tightly coupled to annual cycles in photoperiod, temperature, and energetic allocation, such that baseline female and male transcriptomes are expected to vary substantially between breeding and non-breeding seasons, independent of sex change itself (Laing et al. 2018; Piferrer 2018; Piferrer 2021). This seasonal priming likely shapes the molecular context in which socially induced sex change occurs, resulting in a slower, more progressive transcriptional trajectory characterised by prolonged engagement of immune activation, tissue remodelling, and epigenetic reprogramming pathways (Morais et al. 2012; Budd et al. 2022; Bravo et al. 2023). In particular, post-spawning gonadal regression in seasonal breeders (particularly in females) is commonly associated with elevated inflammatory and immune signalling, which may precondition the gonad for large-scale cellular turnover and thereby facilitate the initiation of sex change outside the peak reproductive period (Morais et al. 2012; Bravo et al. 2023). In contrast, bluehead wrasse inhabit a relatively stable tropical environment and undergo rapid, year-round sex change, consistent with a gonadal transcriptome that is less constrained by seasonal endocrine state and more immediately responsive to social cues, enabling switch-like transcriptional reprogramming (Todd et al. 2018; Todd et al. 2019; Ortega-Recalde et al. 2020). We therefore propose that seasonality acts as an additional regulatory layer in spotty wrasse, modulating the pace and architecture of gonadal reprogramming rather than the ultimate direction of sex change, in line with broader models of layered and environmentally responsive sex-determination systems in vertebrates (Capel 2017; Kitano et al. 2024).

### Conclusions and broader implications

The socially-controlled sex change in the New Zealand spotty wrasse reveals a remarkable degree of phenotypic plasticity underpinned by complex and dynamic gene expression changes across distinct gonadal states. Each phenotypic and transitional stage is defined by a unique transcriptional signature, with classical sex differentiation genes often behaving as expected, but many novel genes emerging as strong candidates for key roles in this process. We describe a process that is likely driven by a flexible regulatory network that integrates both conserved and novel elements to drive one of the most striking transformations in vertebrate biology. These data could have implications for aquaculture, as optimising sex ratios in commercial fish species is frequently a desired outcome. Commercial fish have been routinely treated for decades with androgens or estrogens to induce monosex populations through sex reversal (Pandian and Sheela 1995), for example, trout are gonochoristic, but female trout can be sex reversed to produce XX males using exogenous androgens, which then produce all female offspring when bred with XX females. Understanding how gene expression is regulated in the gonad through the transition may provide insights into master regulators or critical genes and pathways.

## Materials and Methods

### Experimental design and sample collection

Samples were collected during four experimental sampling periods in 2014 (AI14 = Aromatase induced, 2014 study), 2015 (SIP15 = Socially-induced pilot study, 2015), 2016 (SI16 = Socially-induced, 2016 study) and 2018 (SI18 = Socially-induced, 2018 study). Overall, these sampling periods all represent a time series sampling of spotty gonads following (chemically or socially) induced sex change, in different seasons of the year, with timepoints ranging from 21-117 days (total 17 unique timepoints). Two different sampling approaches were used. These were the “all fish from a tank at once” approach, used for experiments in AI14 and SI18, and the “serial sampling” approach, used for experiments SIP15 and SI16. Given the different parameters used in each experiment, sampling timepoints are not always comparable across batches, hence here we compared samples based on histological identification of sex stage. Some samples originally classified as ‘baseline’ females in the SI18 study, these were reclassified as either NBF or BF based on genetic data; this is explored in Supplementary Material section ‘4. Sample metadata exploration’.

### Tissue handling and RNA extraction

Upon sampling, one half of the gonads (one lobe) was flash frozen on a on dry ice/isopentane (C5H12) (Sigma-Aldrich) mixture and stored at – 80 °C for RNA analyses. The other gonadal lobe was preserved for histological analysis (Fig. S2). Briefly, RNA was extracted using Direct-zol RNA kits (Zymo Research), cleaned up when required using RNA clean and concentrator-25 kit with DNase treatment (Zymo Research) and quality assessed using gel electrophoresis, Qubit 2.0 and Fragment Analyzer (Advanced Analytical Technologies Inc.) or Bioanalyzer 2100 (Agilent). For details, see previously published papers (Goikoetxea et al. 2021; Muncaster et al. 2023).

### Reference genome generation

An adult male spotty wrasse was obtained off the coast of North-East New Zealand, Tauranga, between Waihi Beach on the mainland and Mayor Island/Tuhua, and stored at the Toi Ohomai Institute of Technology. From this individual, a liver sample was processed for high-quality genome sequencing. Sequencing was conducted at the Vertebrate Genomes Laboratory at the Rockefeller University, following the Vertebrate Genomes Project (VGP) pipeline v1.6 (Rhie et al. 2021). Briefly, PacBio continuous long reads (CLR) were generated at 71.06x coverage. The CLR reads were assembled into contigs using FALCON-Unzip 2019 Jan-23patch1; false haplotype duplications were removed from the primary haplotype using purge dups v; 10X Genomics linked reads at similar coverage were used to scaffold the contigs using scaff10x v4.1; Bionano optical maps were generated using the DLS enzyme used to further scaffold the assembly using Bionano Solve software; Illumina NovaSeq sequenced Hi-C data generated using the Arima Genomics v1 kit, were used to make a final chromosomal level assembly using Salsa Hi-C v2.2. Polishing of long reads for base call accuracy and gap filling was done using PacBio SMRT link 6.0.0.47841. Further polishing was done after longranger alignment of the 10X short reads, using freebayes v1.3.1. Manual curation [PMID: 33420778] to detect remaining contamination, correct structural errors, remove haplotypic duplications and identify and name chromosomes was performed by the Tree of Life Programme at the Sanger Institute. In the initial automated assembly no contamination was detected, however 18.7Mb of trailing Ns left behind by the Solve were removed. The assembly required 341 manual breaks, 405 manual joins and 9 removals of haplotypic duplication, resulting in an 8% reduction in assembly size (-74.5Mb), a 28% reduction in scaffold numbers and an 11% increase in scaffold N50 with 97.6% of the curated assembly assigned to 24 chromosomes. The genome was deposited into NCBI under accession number GCF_009762535 and assembly ID fNotCel1.pri, further decontamination performed and annotated using the NCBI Eukaryotic Genome Annotation Pipeline (EGAP). The alternate haplotype contigs are under GCA_009762545.1. All assembled and raw data are under the BioProject ID PRJNA562225.

### RNA Sequencing, read trimming, mapping and annotation

RNA samples were sequenced using Illumina TruSeq stranded mRNA library prep with 2x125bp paired end chemistry on Illumina HiSeq 2500 at the Otago Genomics Facility. Raw reads were trimmed using Cutadapt v1.16 (Martin 2011) to retain reads with a quality score PHRED >5 and min length > 40 bp and were deposited into NCBI SRA under BioProject PRJNA631152. Reads were then quality assessed using FastQC v 0.12.1, results aggregated using multiqc v1.24.1, then aligned to the *Notolabrus celidotus* genome (GCF_009762535.1_fNotCel1.pri) using STAR v2.7.9a-GCC-11.3.0 (Dobin et al. 2013) (flags used are in Supplementary Material ‘Mapping and count filtering’). Gene coverage was assessed for each sample using a custom python script [(Stewart 2023) accessed April 2023] that calculates the median percentage coverage (using min-max normalisation) in 10% bins across each gene in a sample. This shows the distribution of reads in each sample across a gene from 5’ -> 3’ and can give an indication of RNA degradation when samples show unreasonably 3’ skewed profiles. Gene ontologies were sourced from the Spotty genome GAF file (GCF_009762535.1, gaf-version: 2.2, generated: 2023-12-09).

### Count matrix, dynamic ‘group-based’ filtering and normalisation

A count matrix with all expressed genes was first generated using the STAR ReadsPerGene output file. A filtered count matrix was then generated, removing very low count genes, by only retaining rows (=genes) where at least 2 values (=samples) were greater than or equal to count of 10, to remove very lowly expressed genes. Counts were then further dynamically filtered to remove all genes with counts <15 in ≥50% of samples within any trait group. ‘Group-based’ filtering was performed in this way as the expectation is that there may be genes that are highly trait specific, and this method allows genes to be retained that may be expressed highly in only a few samples but not others, provided these samples are designated as the same trait (*i.e.,* minimum 50% of the samples in the trait group). For analyses described herein, we used a six trait group count matrix (*i.e.,* F=Female, ET=Early-transitioning, MT=Mid-transitioning, LT=Late-transitioning, TPM=Terminal-phase male, IPM=Initial-phase male), whereby “NBF” and “BF” are combined as one group “F=Female”, used for DGE and WGCNA analyses. Further information on how groups were refined, including re-categorising some trait descriptions, are in Supplementary Material ‘4. Sample metadata exploration’. All count matrices were processed using DESeq2 v1.44.0 (Love et al. 2014) and further transformed using variance stabilisation within the DESeq2 package, when required for visualisation.

### Differential gene expression (DGE) analysis

Differential gene expression (DGE) analysis was performed using DESeq2 v1.44.0 in R v4.4.2 (design = ∼ trait) to a significance cut-off of 0.001. Pairwise comparisons were performed with all trait groups against the female trait group. Results for each trait (ET, MT, LT, TPM, IPM) were then split into genes upregulated against females and genes downregulated against females for all groups. Venn diagrams were generated using VennDetail v1.20.0 (Guo and McGregor 2024) and Venn upset diagram was generated using UpSetR v1.4.0 (Gehlenborg 2019). GOseq v1.56.0 (Young et al. 2010) was then used to perform enrichment and depletion analyses on all comparisons using a significance threshold of 0.05. Following GOseq analysis, the top enriched GO terms (*i.e.,* lowest p-value) were plotted against gene ratio for all genes upregulated in each trait.

### Weighted gene co-expression network (WGCNA) analysis

Weighted gene co-expression network analysis was performed in R v4.3.1 using WGCNA v1.72-5 (Langfelder and Horvath 2008; Langfelder and Horvath 2012). Genes were filtered to remove outliers using `WGCNA::goodSamplesGenes()`, then counts processed using DESeq2 v1.42.1 (design = ∼ 1) and further dynamically filtered (see methods above on dynamic filtering). A soft threshold was picked using a signed network type and visualised using scale free topology model and the mean connectivity.

### Module eigengenes

Modules were generated using `WGCNA::blockwiseModules()`, `TOMType=signed`, `power=20` and `mergeCutHeight=0.25`. Module eigengenes were extracted from the blockwise modules network. Modules were then related to trait groups (F, ET, MT, LT, TPM and IPM) and module-trait association and module membership measures were calculated using `cor()` and `WGCNA::corPvalueStudent()`. Gene expression within modules and across traits were then visualised using violin plots of variance stabilised DESeq2 counts in ggplot2 v3.5.1. GOSeq v1.54.0 was then used to analyse enriched and depleted GO terms within modules (P-value threshold=0.05). Top 50 enriched GO terms were plotted using ggplot2 v3.5.1 by selecting GO terms with the 50 lowest P-values. Gene ratios were calculated from numDEInCat/numInCat, where a gene ratio of 1 indicates that all genes annotated with the GO term (number in category) are differentially expressed (or present) in this gene set (number DE in category).

### Gene-trait correlation

Genes significantly correlated with each trait were calculated as above, using `cor()` and `WGCNA::corPvalueStudent()` and gene expression of the top 30 genes (highest and lowest correlation values) across traits were visualised using box plots of variance stabilised DESeq2 counts in ggplot2 v3.5.1. GOSeq was used to analyse enriched and depleted GO terms in positive and negative gene correlations with traits (P-value threshold=0.05) and the top 5 enriched GO terms were visualised as described above. Statistical analysis was performed using ANOVA between trait groups, using `aov()` in R v4.4.2.

### Classic and putative sex-change genes

Previous research on sex change and sex determination across many species have shown that there is a common suite of ‘classic’ male or female biased genes (Herpin and Schartl 2011; Todd et al. 2016; Capel 2017; Ortega-Recalde et al. 2020). Genes such as *sox9, dmrt1,* and *amh* are typically more highly expressed in males, and genes such as *foxl2, cyp19a1a,* and *bmp15* are typically more highly expressed in females. Other genes implicated in sex change and sex reversal include genes that are important for stem cell pluripotency, cell fate and remodelling, and epigenetic reprogramming. Table 1 shows the list of candidate sex-change genes with notes on expression biases and relevant references. Candidate gene expression was plotted using ggplot2 boxplot and statistical difference in gene expression between female and TP male trait groups was performed using ANOVA.

## Acknowledgments

We would like to thank Jodi Thomas and Erin Damsteegt for their technical assistance in preparing RNA samples for library preparation and sequencing, Bettina Haase for their role in Pacbio, 10X genomics, and Hi-C Sequencing, Jacquelyn Mountcastle for their role in DNA extraction and processing for Bionano, Chai Fungtammasan for their role in assisting with genome assembly, Arang Rhie for guidance on the assemblies, and Ying Sims for setting up the gEVAL data set.

## Author Contributions

A.G., S.M., E.T., O.O-R. and N.G. conceived and designed the original studies;

A.G., S.M., O.O-R. and E.T. carried out fish experimentation and generated RNA-seq samples;

O.F., E.J., A.T. and K.H. performed genome generation, assembly and annotation;

C.V.D.B. and K.K. analysed transcriptomic data;

C.V.D.B. wrote the manuscript;

C.V.D.B., K.K., T.H., E.J., E.T., K. H., A.G., and N.G. edited and revised the manuscript. All authors read and approved final manuscript.

## Funding

This research was made possible by Royal Society Te Apārangi Marsden funding to Neil Gemmell (Grant no. UOO1308 and UOO2115) and by Howard Hughes Medical Institute funding to Erich Jarvis.

## Conflict of interest

The authors declare no competing interests.

## Data availability

The genome assembled in this manuscript is available at NCBI Genome Datasets under the accession GCF_009762535.1 (Genome assembly fNotCel1.pri). The RNA-seq data analysed in this study have been deposited in the NCBI SRA archive with the accession PRJNA631152.

## Notes

### Competing Interest Statement

The authors have declared no competing interest.

