## Supplementary Material for "Gonadal remodelling in socially driven sex change is associated with novel and known sex genes, epigenetic reprogramming, and inflammation-mediated apoptosis"

### 1. Spotty phenotypes and gonad histology

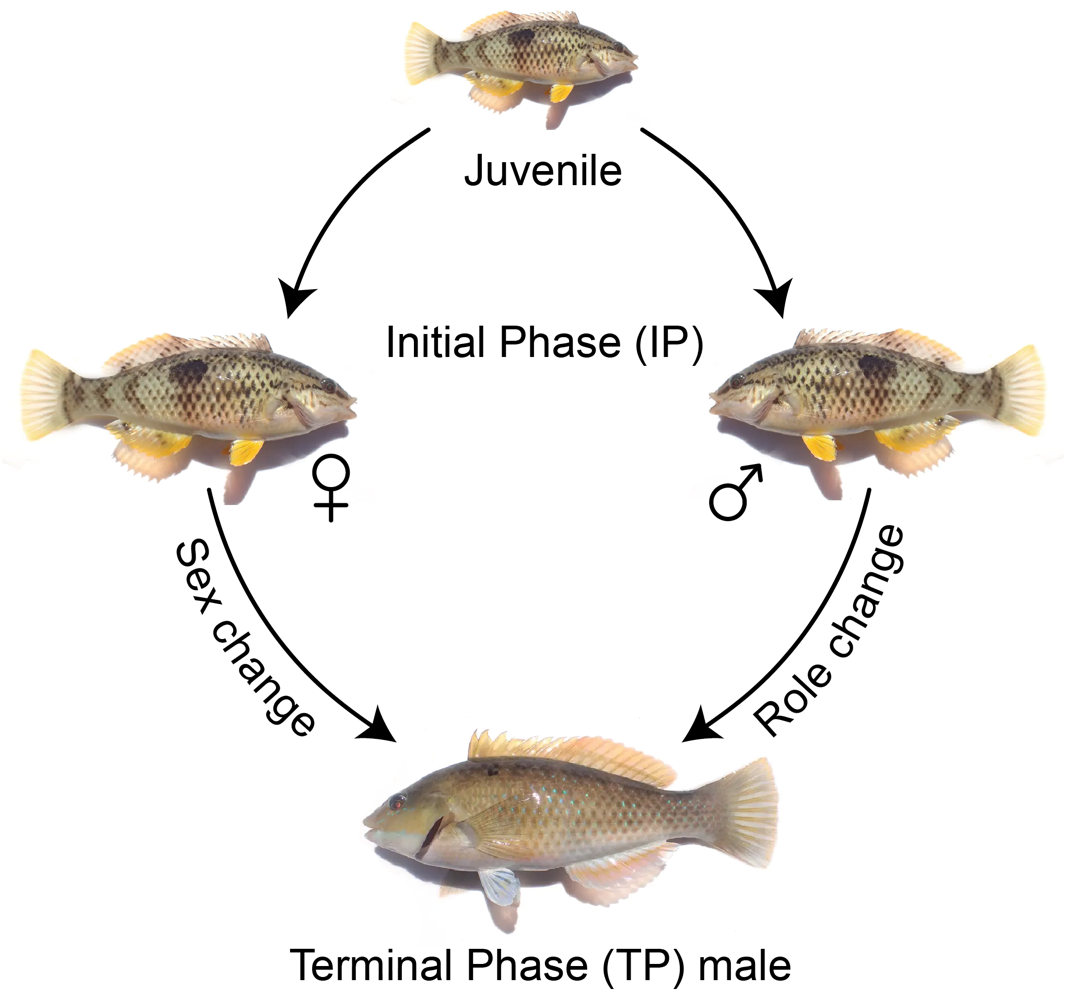

**Figure S1**. Sex phenotypes in spotty wrasse. Juvenile spotty differentiate as either Initial phase females or males, which are virtually indistinguishable. Both IP phenotypes can change to the Terminal phase male, which has distinct colouration and marking differences. Images copyright of Ryota Hasagewa, 2023, adapted with permission. Original images can be found here: <https://zukan.com/fish/internal4301> ; <https://zukan.com/fish/leaf165371>

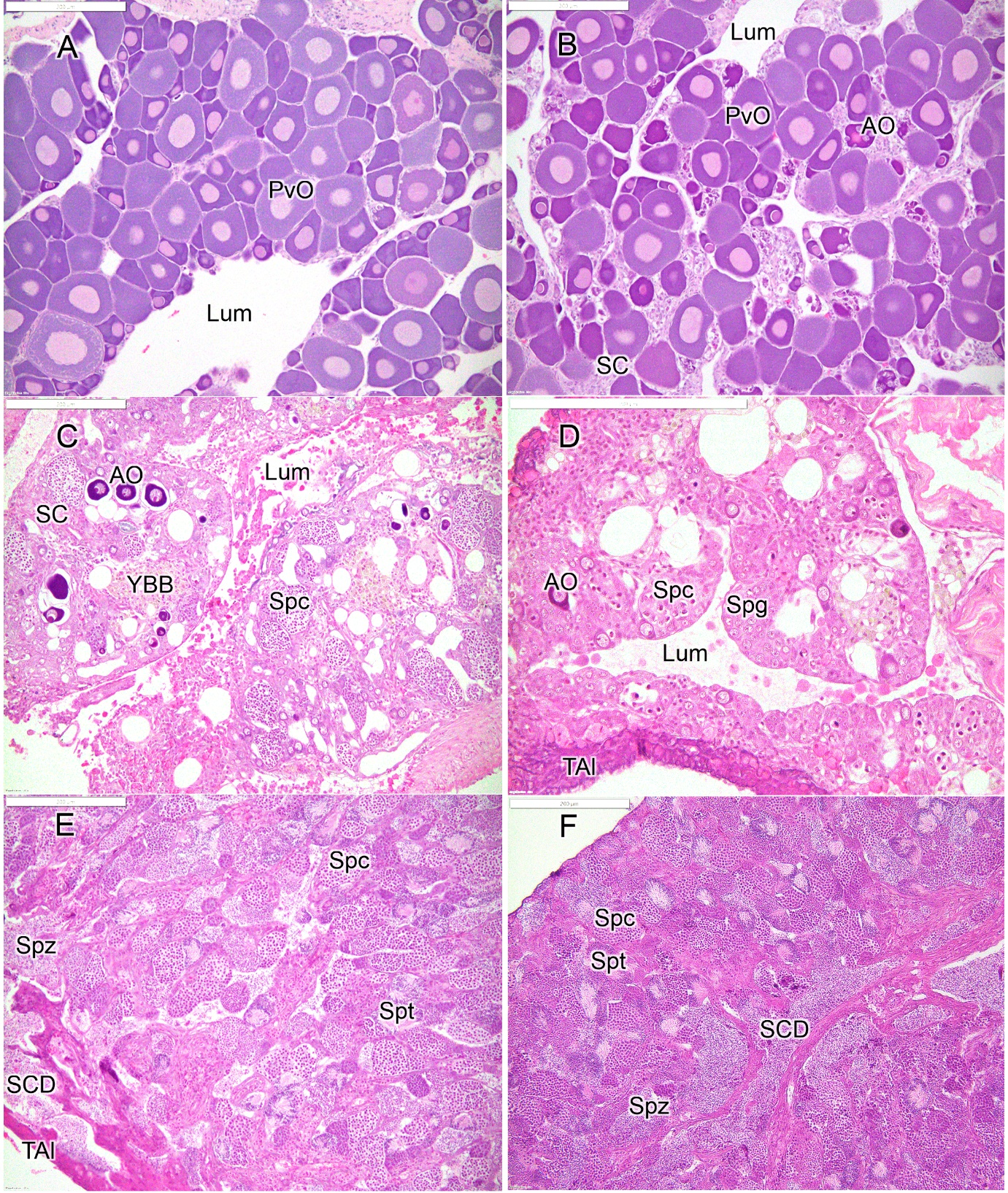

**Figure S2.** Histological stages of gonadal development in spotty wrasse: A) initial phase female; B) early transitioning fish; C) mid transitioning fish; D) late transitioning fish; E) terminal phase male; F) initial phase male. Abbreviations: atretic oocyte (AO), gonadal lumen (Lum), previtellogenic oocyte (PvO), stromal cells (SC), sperm collecting duct (SCD), spermatocytes (Spc), spermatogonia (Spg), spermatid (Spt), spermatozoa (Spz), tunica albuginea (TAl), yellow-brown body (YBB). Scale bars = 200 µm.

### 2. RNAseq datasets

#### *2a. Experimental design and sample collection*

Most samples (2014, 2016 and 2018, but not 2015) were collected as part of work published by (Goikoetxea et al. 2021) and (Muncaster et al. 2023). However, while gonads were collected and RNA extracted for gene-of-interest analyses in these studies, full transcriptome analysis was not performed at that time.

In the AI2014 study, sex change was induced using the aromatase inhibitor fadrozole, between August and September (Winter-Spring). Social sex change was blocked in these fish by placing a TP male in each tank. All fish from a single tank were sampled on a given day. This study has been published by coauthors (Goikoetxea et al. 2021).

In the SIP2015 study, a pilot study of social induction of sex change was performed within breeding season (September-December; Spring-Summer), by removing TP males from the tank. The largest IP fish from each tank was terminally sampled on day 43 (n=6), day 70 (n=6) and day 84 (n=6). Not all samples were used for RNAseq (see Supplementary Table S1). Fish were sampled serially from each tank, with all tanks being sampled on a given day. The fish used in this study have not been previously published elsewhere.

In the SI2016 study, social induction of sex change was performed within breeding season (September-December; Spring-Summer), by removing TP males from tanks. Control tanks were also set up where TP males were left in the tank. The largest IP fish from each tank was terminally sampled on days 0, 30, 50, 60, 65, or 66 (end of experiment) (n total = 10 per sampling day). Fish were sampled serially from each tank, with all tanks being sampled on a given day. Not all samples were used for RNAseq (see Supplementary Table S1 for sample list for this study). This study has been published by coauthors (Goikoetxea et al. 2021).

In the SI2018 study, social induction of sex change was performed outside breeding season (January-April; Summer-Autumn), by removing TP males from tanks. Fish were sampled over a time series as follows; day 1 (n = 5), day 11 (n = 5), day 26 (n = 10), day 36 (n = 10), day 55 (n = 10), day 92 (n = 9). A further five IP females were terminally sampled on day 0 to provide a baseline indication of reproductive status (BLF; baseline female). A total of 49 samples were used for RNAseq, including all 5 baseline females (day 0) and 1 TP male. All fish from a single tank were sampled on a given day. This study has been published by coauthors (Goikoetxea et al. 2021; Muncaster et al. 2023).

Given the different parameters used, sampling timepoints were often not comparable across batches. This is because for some tanks, the most dominant fish was only removed once (‘all fish from a tank at once’ approach) and for other tanks, the most dominant fish changed through the experiment (‘serial sampling’ approach). As the removal of the most dominant fish can induce sex change, regardless of what sex the fish is (Quertermous et al. 2025), this in effect may ‘reset’ the clock for that tank. Hence herein we compared samples based on histological identification of sex stage only, rather than by timepoint, which allowed us to compare gene expression based on the known phenotype of the sample, rather than the assumed level of sex change based on time.

### 3. Quality assessment, mapping and count filtering

#### 3a. FastQC/multiqc reports

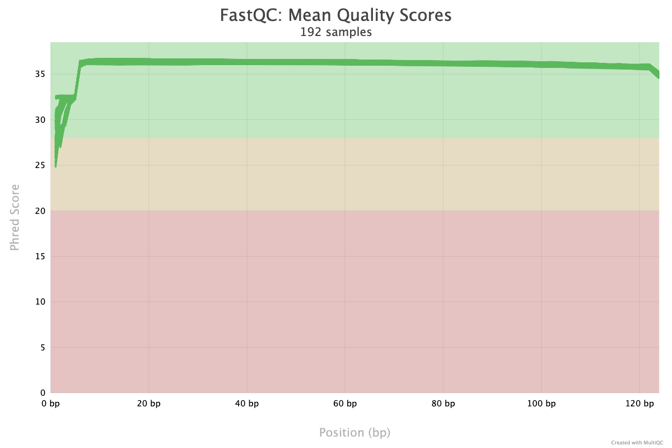

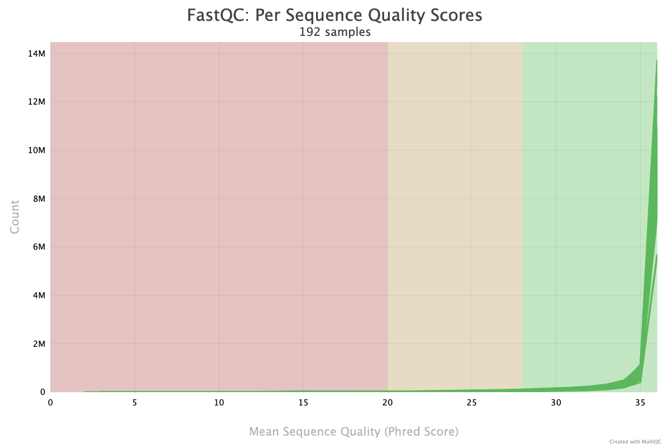

**Figure S3.** Multiqc-aggregated fastQC report results showing > Q30 overall sequence quality for 192 files (R1 and R2 paired-end read files for 96 samples).

#### 3b. STAR alignment flags

```
STAR \
  --genomeDir /path/to/genomeDir \
  --readFilesIn ${INPUT_FILE_1} ${INPUT_FILE_2} \
  --outFileNamePrefix ${OUTPUT_FILE} \
  --outSAMtype BAM SortedByCoordinate \
  --quantMode TranscriptomeSAM GeneCounts \
  --genomeSAindexNbases 13 \
  --sjdbGTFfile GCF_009762535.1_fNotCel1.pri_genomic.gtf \
  --readFilesCommand zcat \
  --sjdbOverhang 100 \
  --twopassMode Basic \
  --outSAMattrIHstart 1 \
  --outSAMattributes NH HI AS nM ch \
  --outSAMprimaryFlag OneBestScore \
  --outSAMmapqUnique 60
```

#### 3c. Mapping results

Samples were further investigated to visualise sequencing quality. Fig. S4a shows a relatively high percentage of uniquely mapped reads across all samples (as determined by STAR), with a slightly lower average across SI18 samples. An average of 86.15% of all reads uniquely mapped to the genome (range of 72.81 % - 91.06 %). Fig S4b shows how reads map across all genes, as a function of percentage coverage of the gene in 10% bins from 5’->3’. As expected with polyA capture, there is an upwards trend towards the 3’ end, indicating a higher coverage at the 3’ end of genes. Some samples, and particular evident in sample SI18_38G (MT), show a very high 3’ bias, with poor coverage across the 5’ end, indicating poorer quality of these reads (i.e., likely more degraded mRNA).

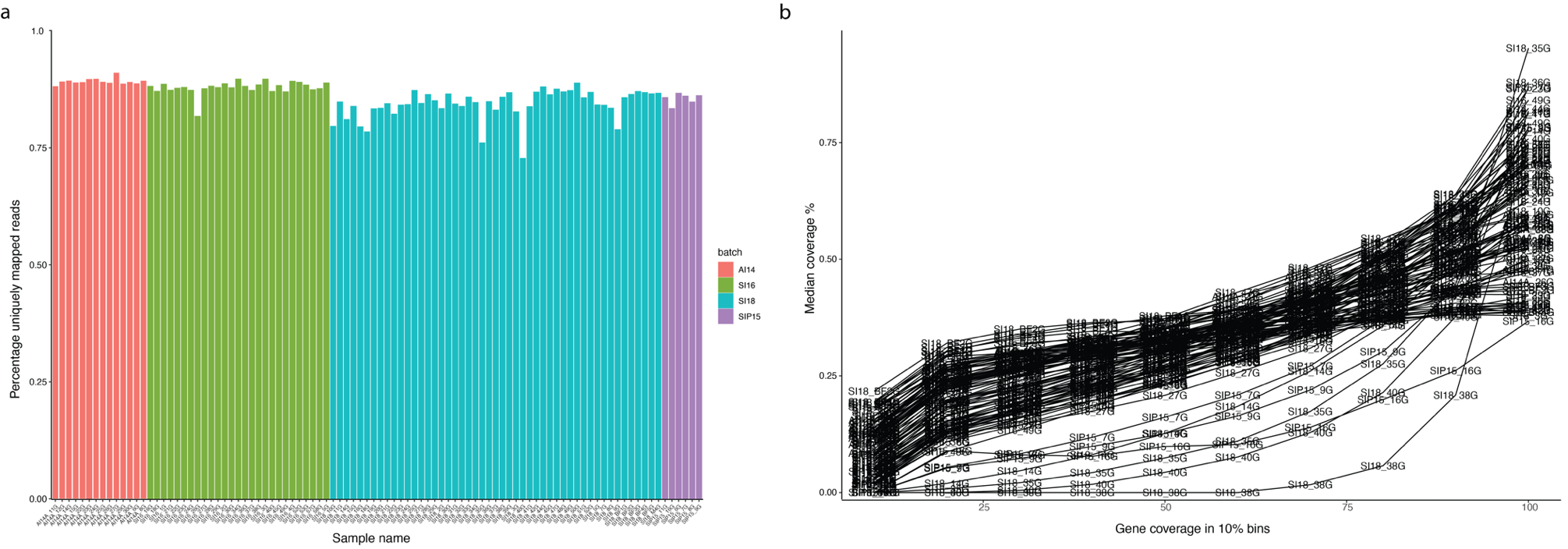

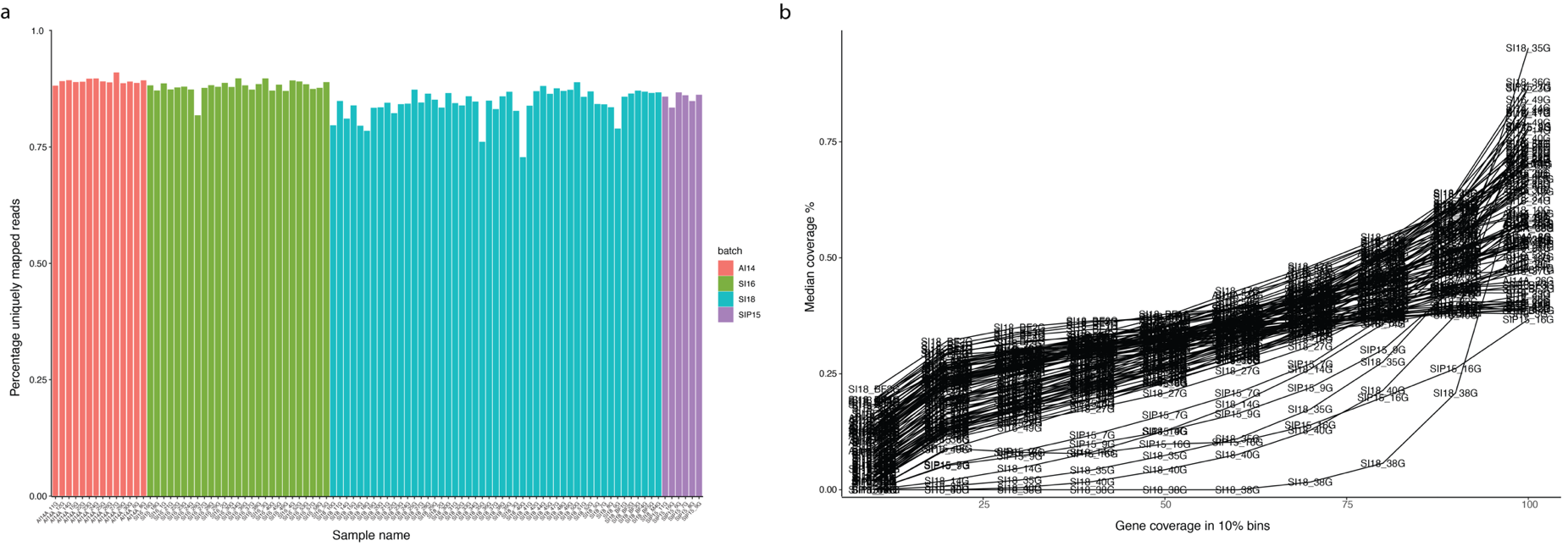

**Figure S4.** Mapping results. A) barplot showing percentage of uniquely mapped reads for all 96 samples. B) Gene coverage QC results for all 96 samples. Gene coverage is calculated in 10% bins (5’ -> 3’ across the x-axis) and is averaged across all expressed genes for each samples. Median coverage percentage is calculated using the min-max normalised coverage.

#### 3d. Group filtering

After count matrices were generated from STAR, very low counts were first filtered out, then a dynamic ‘group filtering’ approach was employed to further filter count matrixes, rather than using a strict count cut-off. This approach was chosen as it was expected that there may be genes that are highly trait specific, and this method allows genes to be retained that may be expressed highly in only a few samples but not in any others, provided these samples are designated as the same trait (i.e., minimum 50% of the samples in the trait group). As this method therefore is dependent on how many groups are used, two matrices were generated. Figure S5 shows the average counts for the genes included and excluded from analysis across each trait group, using 7 trait groups (Fig. S5a) and 6 trait groups (Fig S5b). The number of genes retained in each was very similar and made very little difference to any downstream analyses.

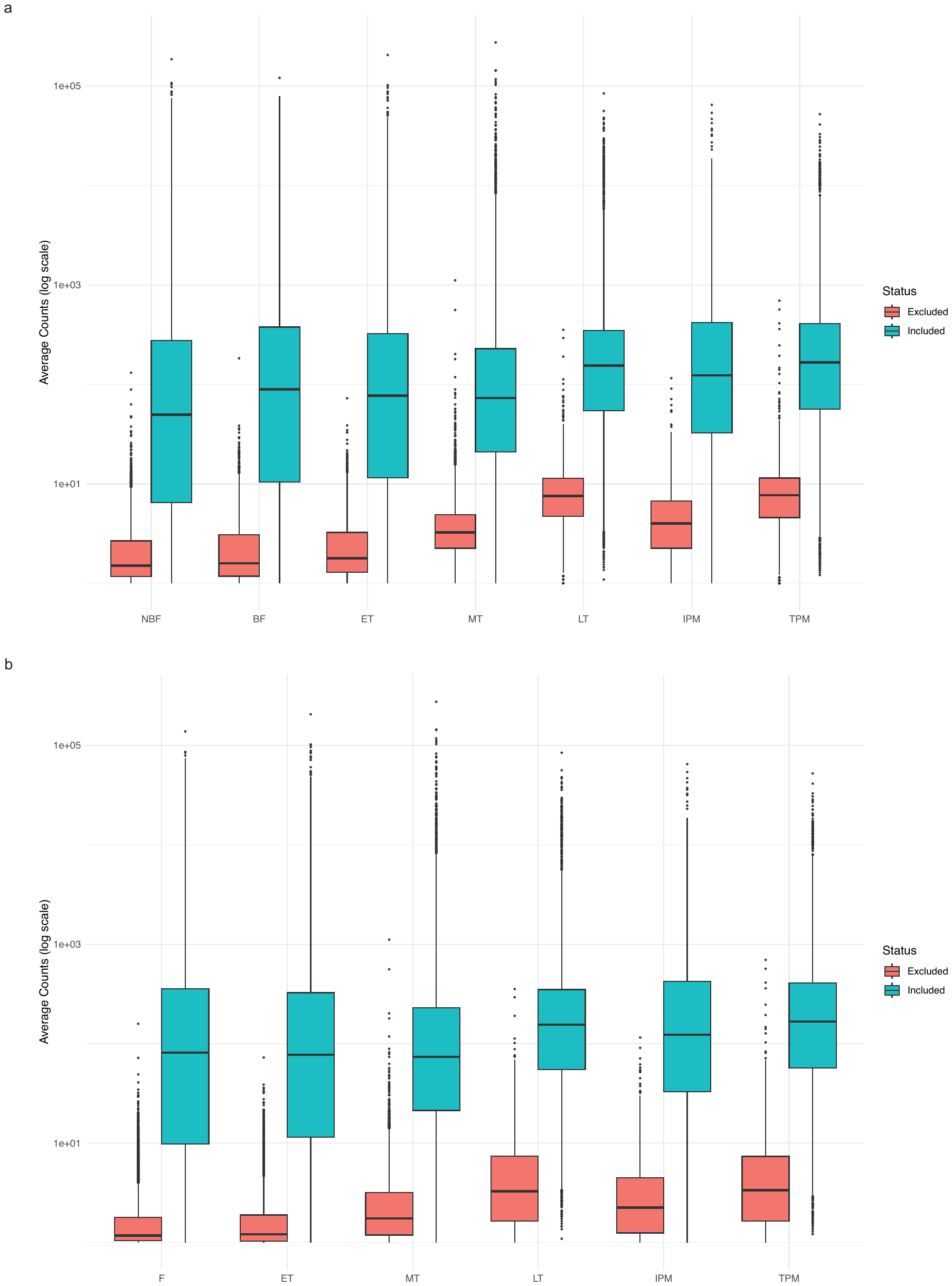

**Figure S5.** Average counts of genes included and excluded from each trait group after filtering. a) group filtered using 7 trait groups (i.e., female category split into NBF=non-breeding female and BF=breeding female), included genes = 18,374 total; b) group filtered using 6 trait groups (i.e., female single category), included genes = 18,320 total.

### 4. Sample metadata exploration

#### *4a. Initial assessment using PCA*

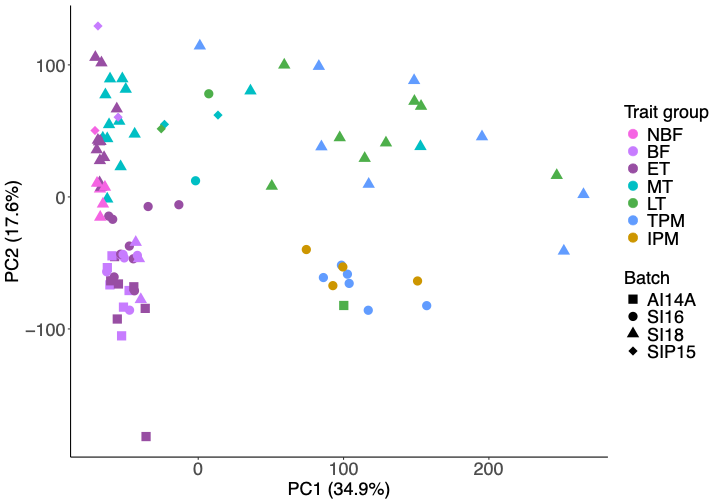

**Figure S6**. PCA plot using DESeq2 filtered counts (low counts removed, 22,231 genes remaining) showing variance distribution by trait group and batch.

#### *4b. Further exploration and refinement of trait groups*

In the original experimental design for SI18, 5 samples were categorised as “Baseline females” and were sampled on day 0 of the experiment. No histology was performed for these samples. However, initial PCA analysis of these samples (figure S7) showed SI18_BF1G, SI18_BF2G and SI18_BF3G clustered with BF, and samples SI18_BF4G and SI18_BF5G clustered with NBF. These were therefore reclassified as such for some investigatory analyses. However, as this was not determined histologically, and given the overlap genetically in female states (as well as overlap with ET) seen in figure S8, for most analyses described in this study (i.e., DGE, WGCNA) all BLF, NBF and BF samples were grouped together as “female” for comparisons between states.

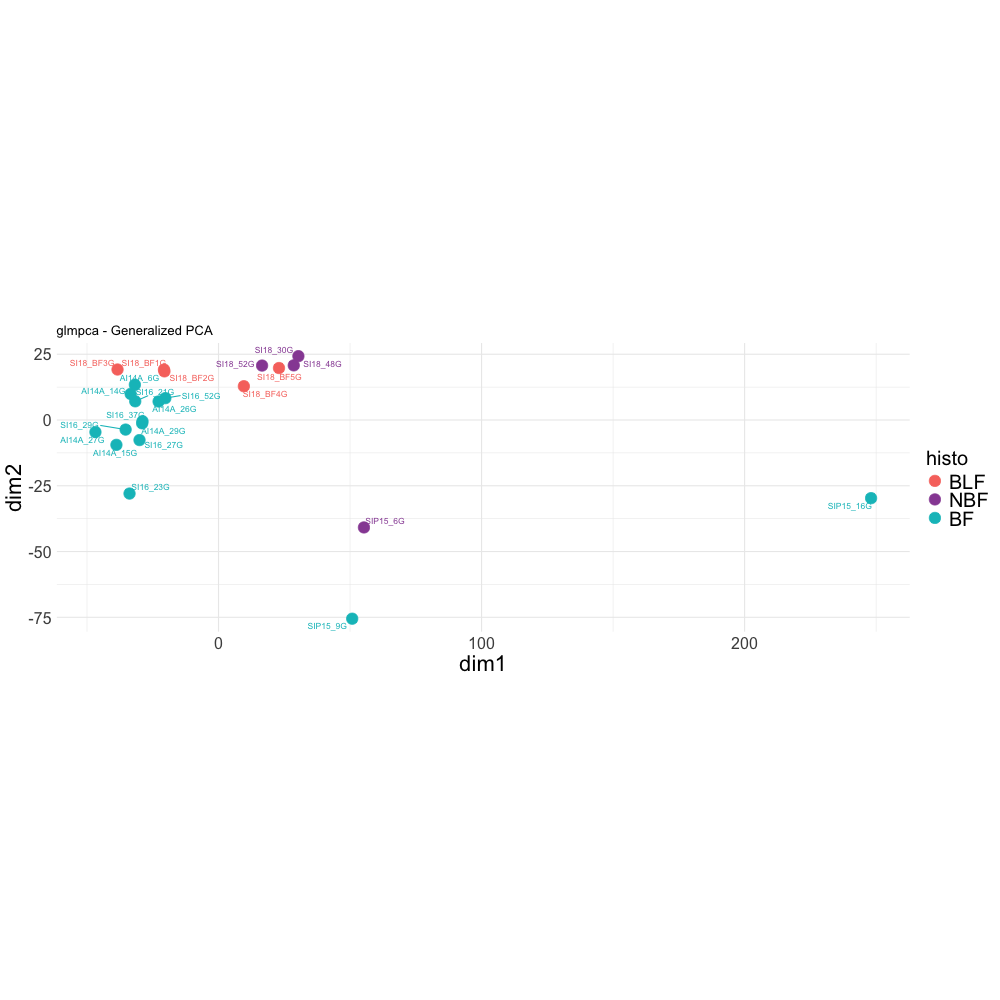

**Figure S7.** GLM PCA using only samples designated as BLF (baseline female), NBF (non-breeding female) or BF (breeding female). Counts were processed using DESeq2 and low counts removed (dds <- dds[rowSums(counts(dds) >= 15) >= 17,]).

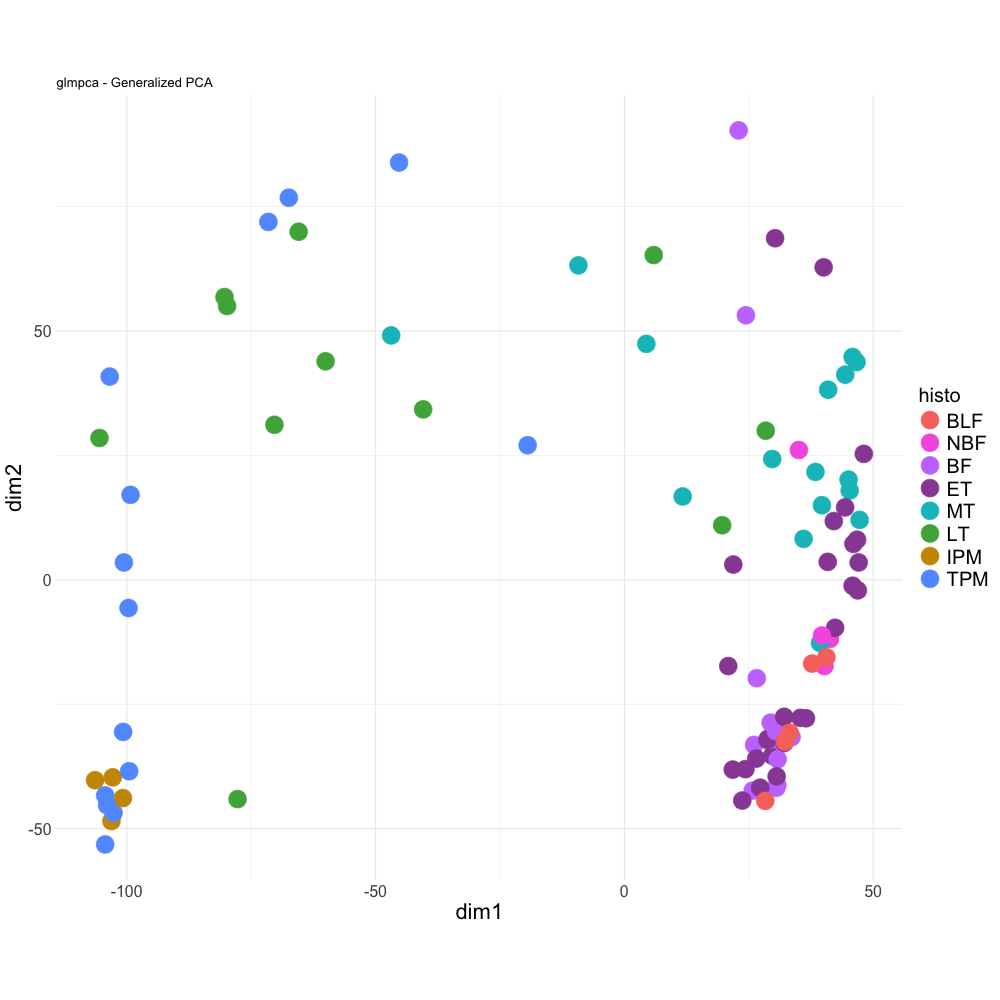

**Figure S8.** GLM PCA using all samples (in comparison to the figure above, where only female samples are shown). Overlap can be seen between BLF, BF, ET and NBF samples. Counts were processed using DESeq2 and low counts removed (dds <- dds[rowSums(counts(dds) >= 15) >= 72,]).

*Exploration of gene count changes over sex change*

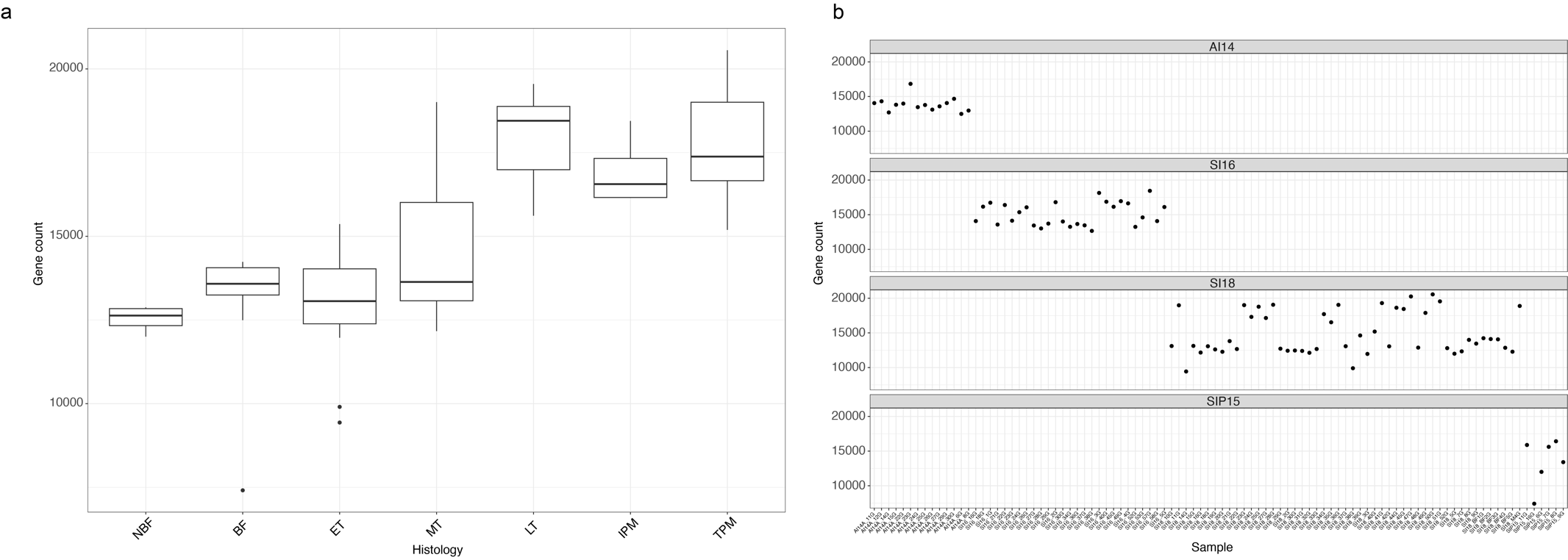

**Figure S9.** Number of expressed genes (not expression levels) in each trait group. A gene was only counted as expressed if it had a minimum of 10 counts (STAR counts) for the given sample. This plot does not show expression levels for each gene. There is a clear trend showing that the number of genes that are expressed in each trait group increases through the transition and male gonads express more genes than female gonads.

### 5. DGE

To further expand on Figure 2 in the main manuscript, the gene expression across traits and the top 50 gene ontology terms for genes upregulated and downregulated in each trait (vs F) are visualised here below in Figures S10 and S11. We also explored the expression of the 36 genes uniquely upregulated in all traits vs female, shown in Figure S12.

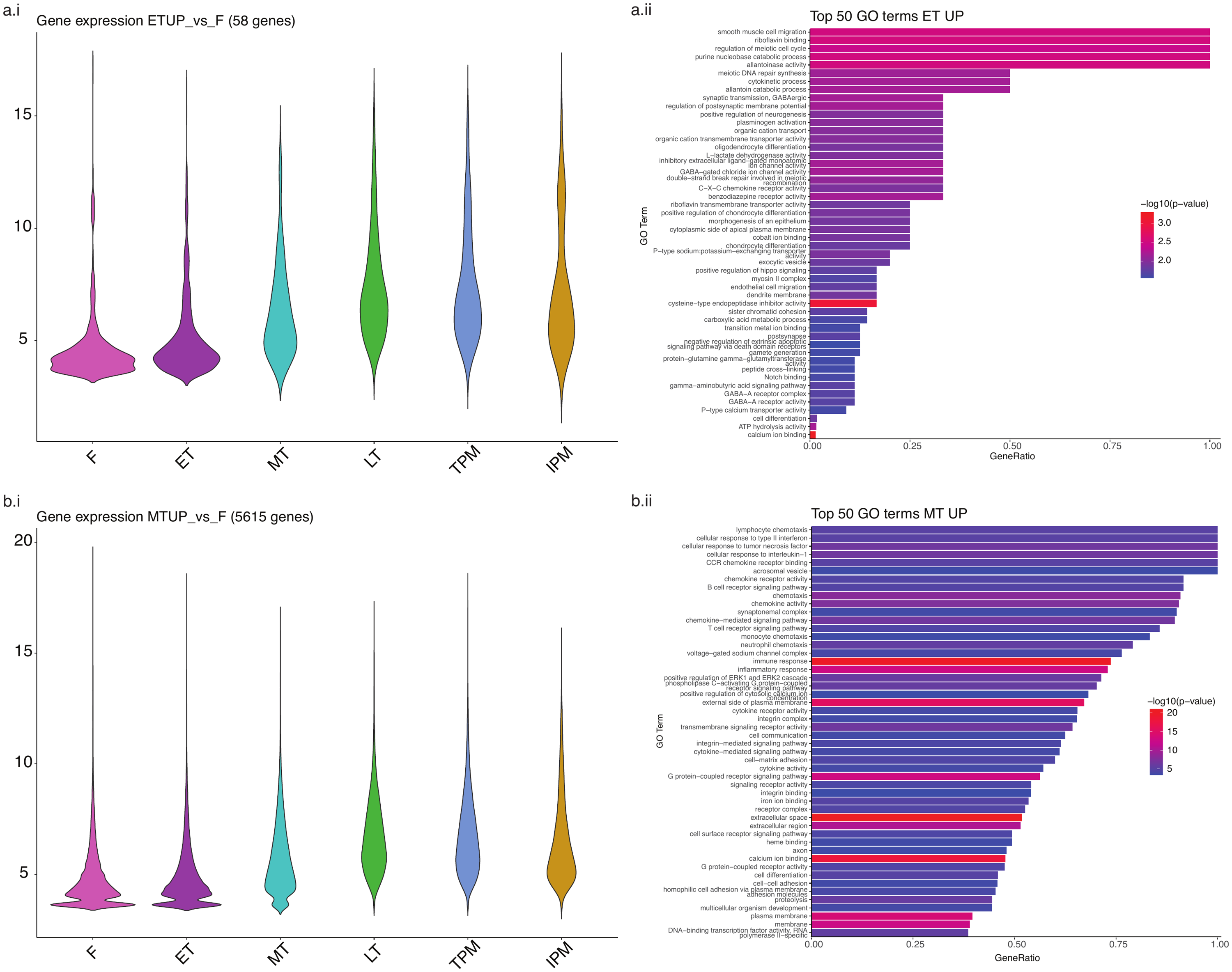

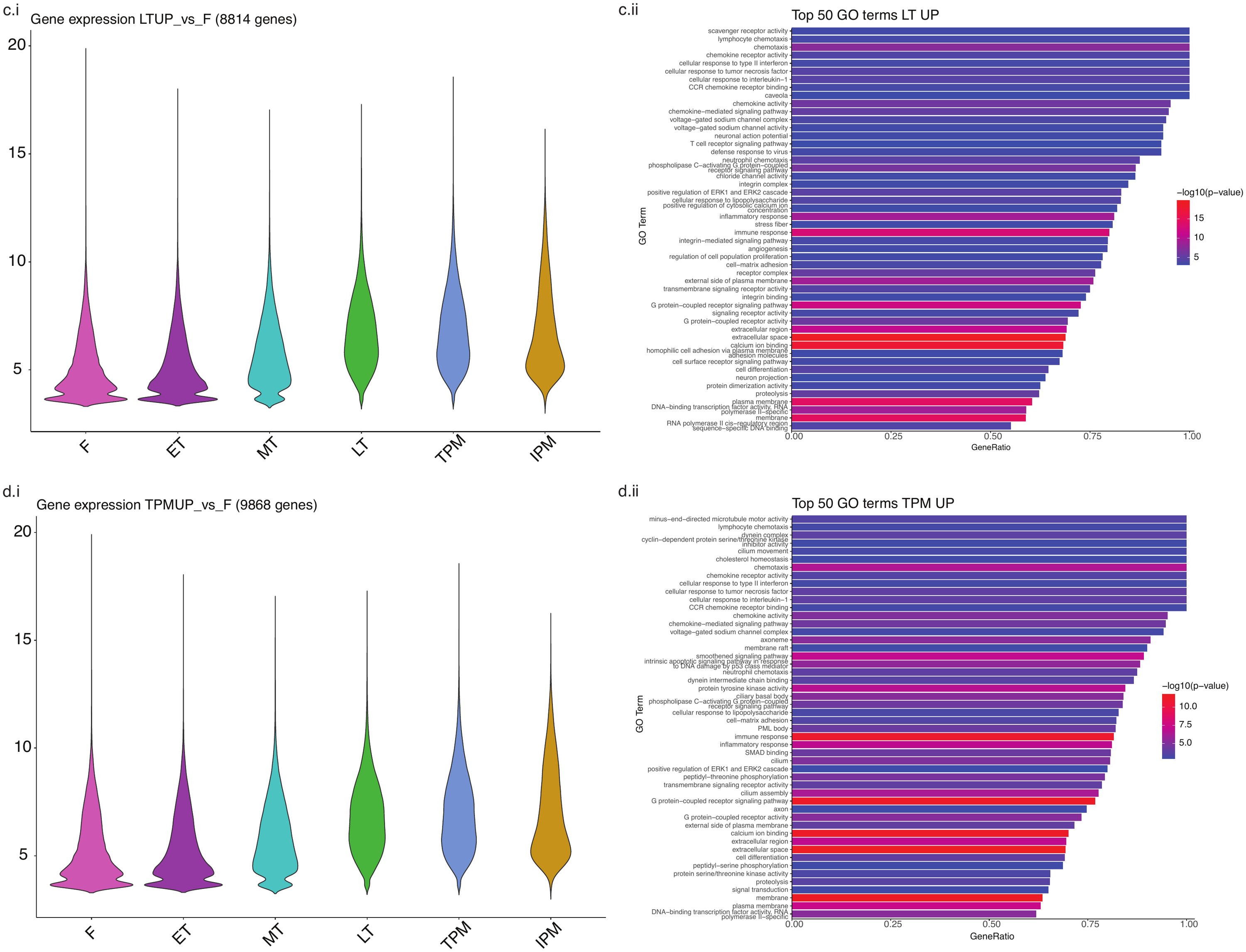

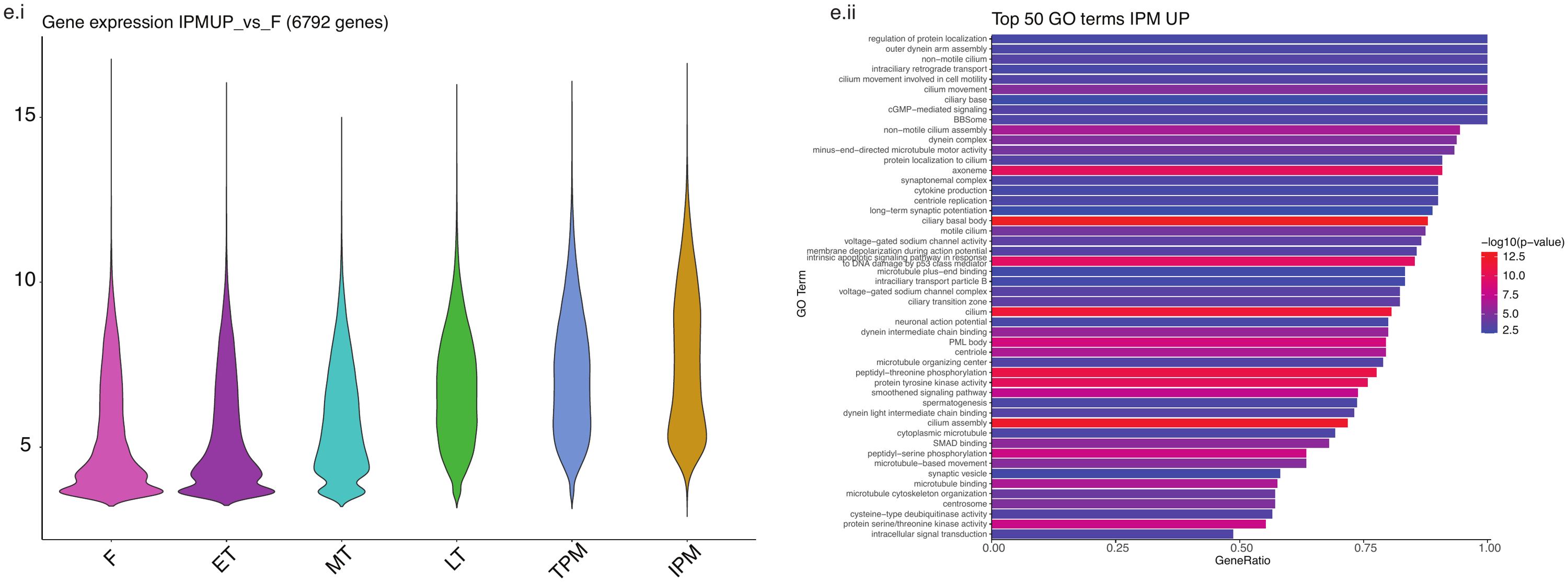

**Figure S10**. **Gene expression and top 50 enriched GO terms for genes upregulated in each trait, vs female**. a-e.i) Gene expression across traits of genes significantly upregulated in X trait, vs F and a-e.ii) top 50 GO terms enriched in genes upregulated in X trait.

Gene expression shown as vst-normalised DESeq2 counts. Gene ratio calculated from numDEInCat/numInCat. See Supplementary Tables S3-S8 for full GOseq data for each trait vs female DGE.

This figure directly expands on Figure 2d-h in the main manuscript, which only shows the top 5 GO terms.

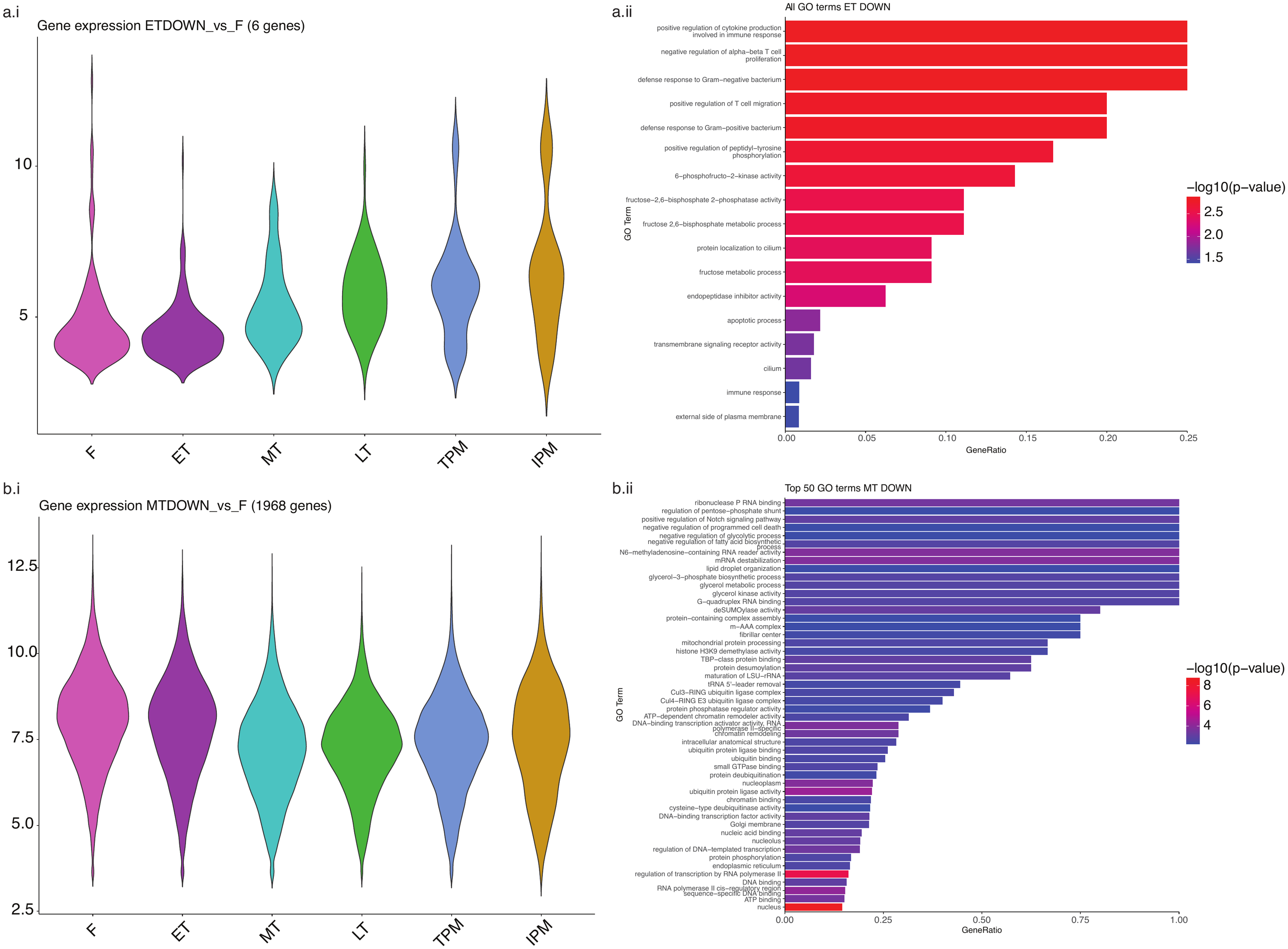

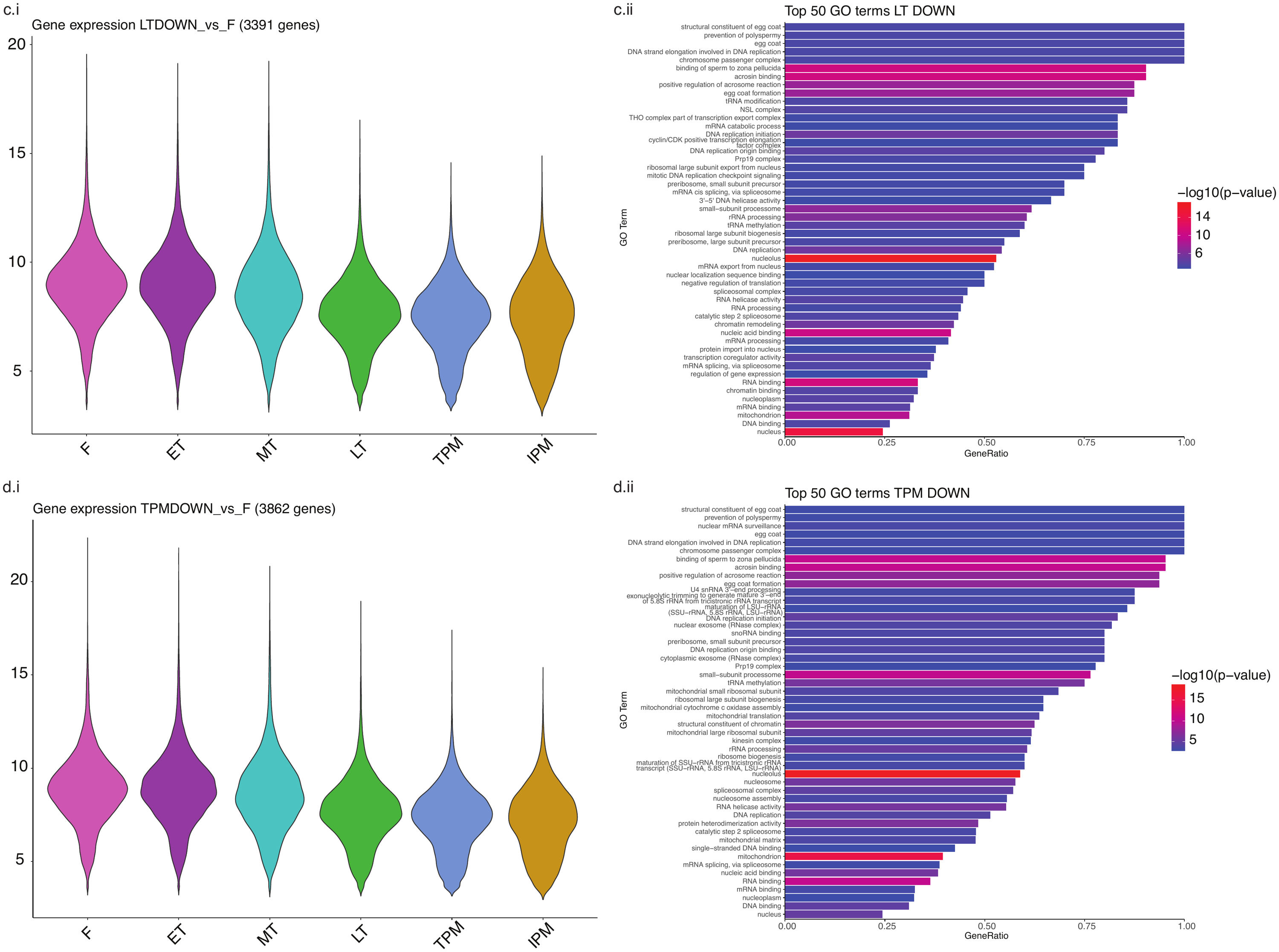

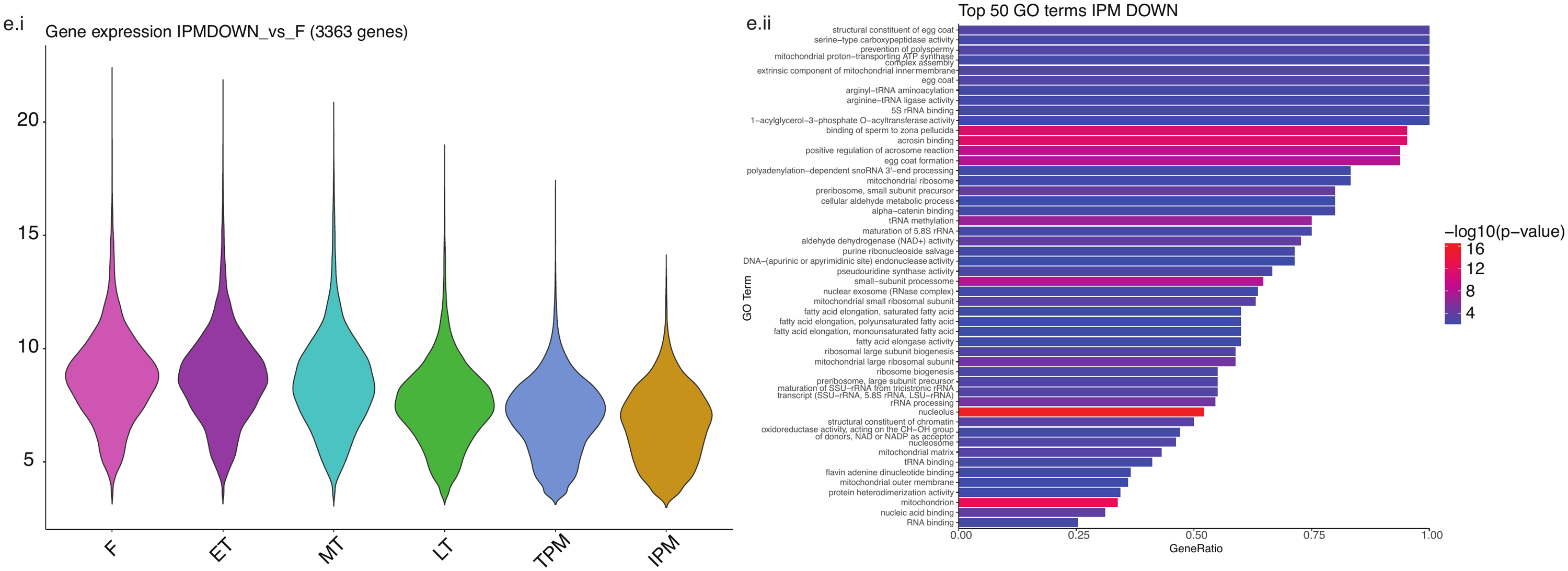

**Figure S11. Gene expression and top 50 enriched GO terms for genes downregulated in each trait, vs female**. a-e.i) Gene expression across traits of genes significantly downregulated in X trait, vs F and a-e.ii) top 50 GO terms enriched in genes downregulated in X trait (for ET, all GO terms enriched in genes downregulated in ET are shown, as there were not 50 GO terms).

Gene expression shown as vst-normalised DESeq2 counts. Gene ratio calculated from numDEInCat/numInCat. See Supplementary Tables S3-S8 for full GOseq data for each trait vs female DGE.

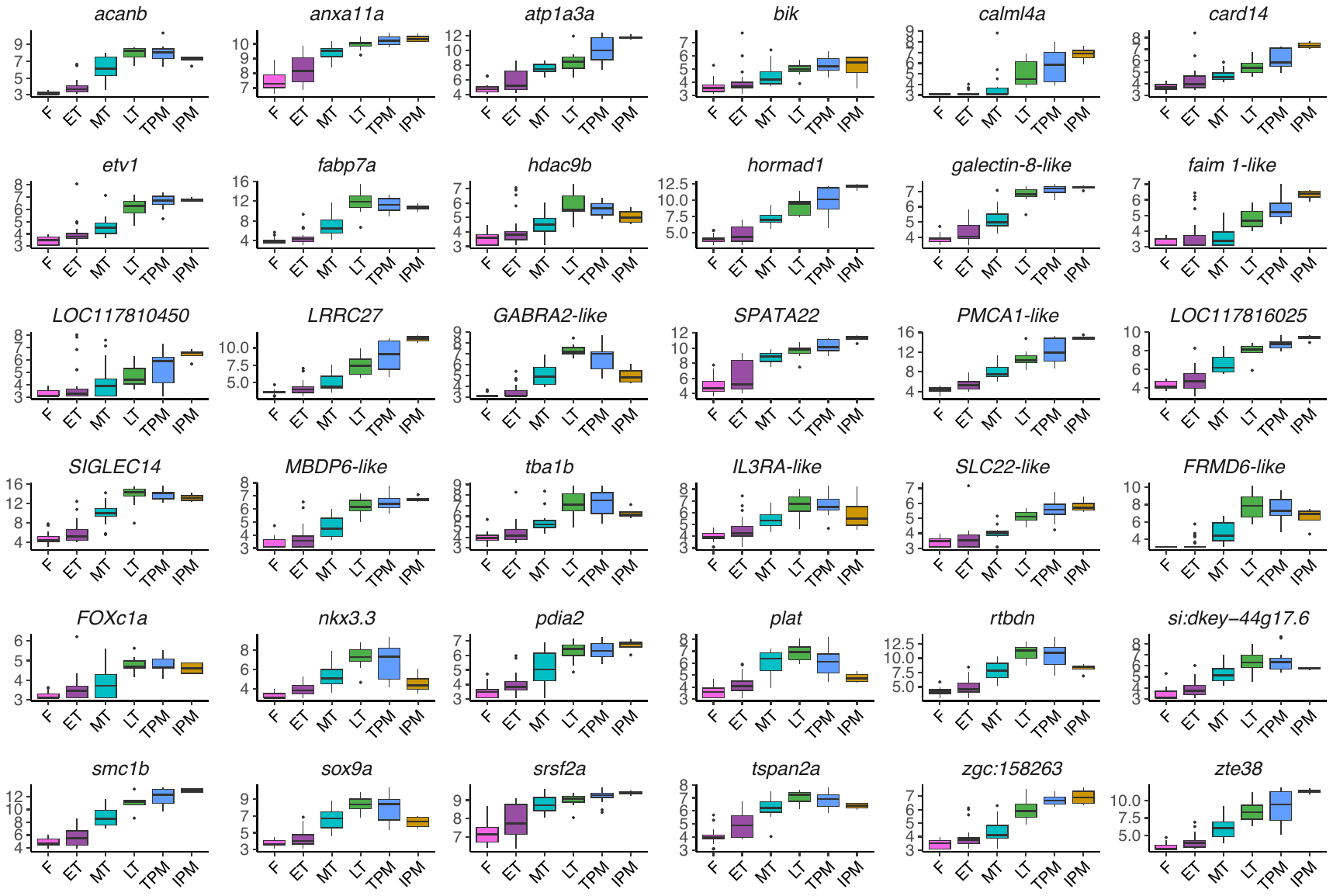

**Figure S12.** Expression across traits of the 36 unique genes that are significantly upregulated in all traits, compared to females (Manuscript Figure 2b). Expression on y-axis shown as variance stabilised DEseq2 counts. Gene list in Supplementary Table S2. Note that for genes where the ID was designated in the genome as “LOC#########”, these were changed here to the best descriptive gene name (e.g., LOC117812621 = GABRA2-like). Two genes (LOC117810450 and LOC117816025) are described only as uncharacterized genes.

*N.B.: In our previous work on spotties (Muncaster et al. 2023), the gene sox9a was found to be downregulated in male gonads, which is opposing to other literature on this gene in vertebrates (Herpin and Schartl 2011; Liu et al. 2015; Todd et al. 2019; Ortega-Recalde et al. 2020; Vining et al. 2021). Here, however, we found sox9a to be upregulated in males, in line with other vertebrate literature.*

### 6. WGCNA

This section describes supplementary analyses for weighted gene correlation network analysis, with data on soft power detection, module gene expression exploration and gene-trait correlation exploration.

To choose a soft power for WGCNA analysis, scale free topology and mean connectivity were both visualised in relation to power in figure S13.

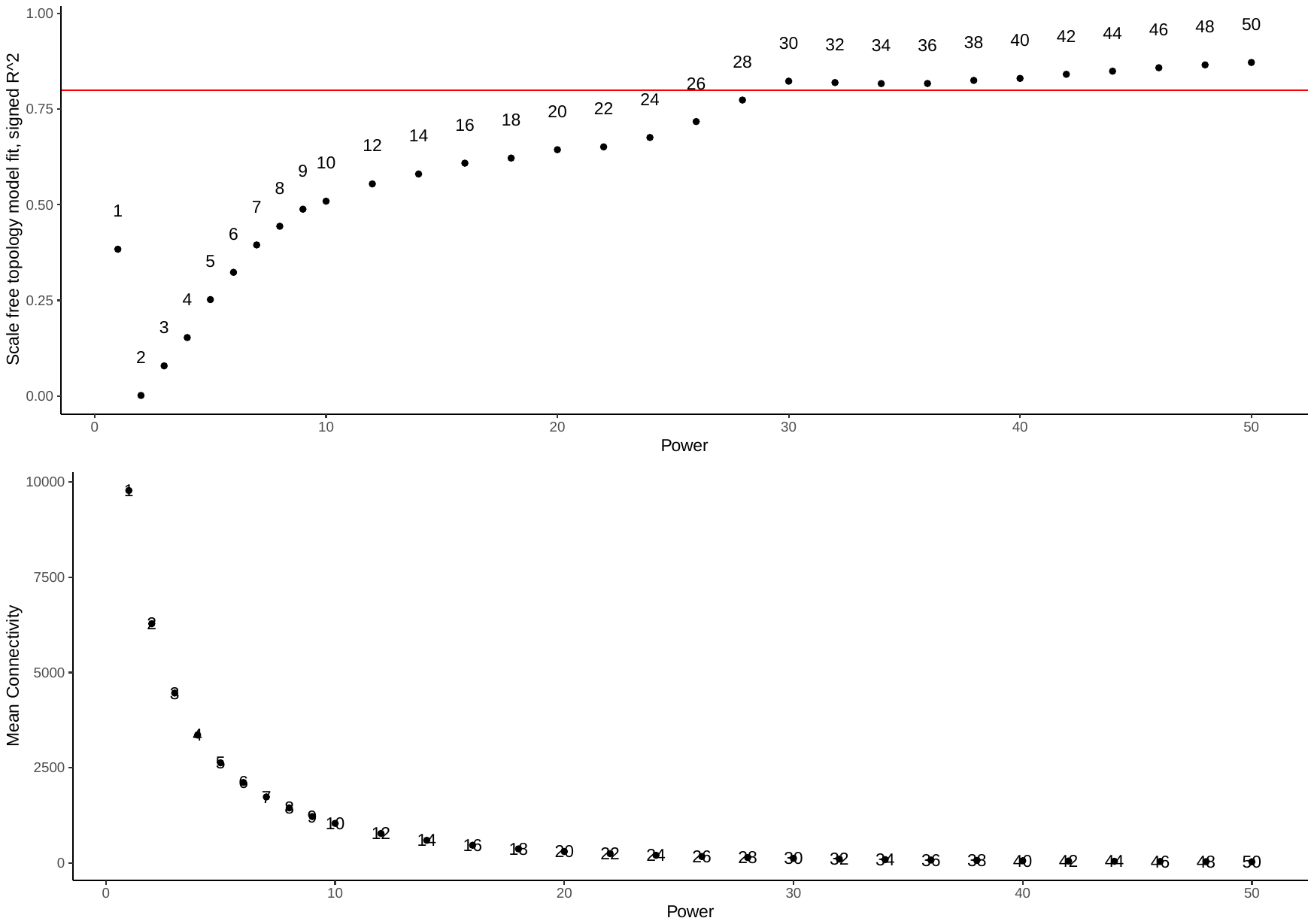

**Figure S13**. Soft power detection and visualisation. a) Scale free topology model. b) Mean connectivity.

#### 6a. WGCNA: modules

Of the nine modules detected (Supplementary Figure S14), two modules (turquoise and blue) show a clear masculinising trend, with strong positive correlation with TP males and simultaneous negative correlation with F and ET traits. Two modules (magenta and yellow) show a clear feminising trend, with strong positive correlation with females and simultaneous negative correlation with MT, LT, TPM and IPM (Figure S14c). The GO terms for these modules are further explored visually (Supplementary Figure S15) and full terms are listed in Supplementary Table S10.

1. *Turquoise – male-biased*

The top GO terms for this module are immune system functions, such as, chemotaxis /chemokine activity and inflammatory response (e.g., MHC class II protein binding, cytokine activity, monocyte/lymphocyte chemotaxis and activity), cell signalling, proliferation and migration (e.g., growth factor, positive regulation of cell population proliferation, positive regulation of cytosolic calcium ion concentration, intermediate filament of cytoskeleton organization, cell adhesion mediated by integrin, G protein-coupled receptor signaling pathway, positive regulation of ERK1 and ERK2 cascade) and transcriptional regulation (e.g., DNA-binding transcription factor activity, RNA polymerase II-specific, RNA polymerase II cis-regulatory region sequence-specific DNA binding) (Fig. S14c; correlation value 0.5).

1. *Blue module – male-biased*

The top GO terms for this module are comprised of primarily spermatogonia growth and proliferation genes. GO terms include: terms directly related to sperm (e.g., sperm midpiece, sperm axoneme assembly, flagellated sperm motility), movement (e.g., motile cilium assembly, cilium-dependent cell motility, cilium movement), mitosis (centriole replication, regulation of mitotic cell cycle, G1/S transition of mitotic cell cycle) and kinase/phosphorylation activity (e.g., protein tyrosine/serine/threonine phosphorylation activity) (Fig. S14c; correlation value 0.61).

2. *Magenta module – female-biased*

The top GO terms for this module are pre-translation processing (e.g., tRNA-intron endonuclease activity, rRNA methyltransferase activity, maturation of 5S rRNA, polynucleotide 3’-phosphatase activity), post-translation protein processing (e.g., Golgi calcium ion homeostasis, protein deglycosylation, protein localisation to secretory granule), sterol/steroid processing (e.g., lanosterol synthase activity, isoprenoid biosynthetic process, cholesterol biosynthetic process, triterpenoid biosynthetic process) and transcription (e.g., TBP-class protein binding, regulation of transcription by RNA polymerase II, RNA polymerase II cis-regulatory region sequence-specific DNA binding) (Fig. S14c; correlation value 0.5).

3. *Yellow module – female biased*

The top GO terms for this module indicate an increase in mitochondrial processing/replication (e.g., mitochondrial fission, mitochondrion, MICOS complex), DNA replication and repair (e.g., DNA replication initiation, DNA replication, DNA clamp unloader activity, DNA replication origin binding, DNA repair, MCM complex), enzymes and metabolic process (e.g., glucosidase II complex, 3-hydroxyacl-CoA dehydratase activity, endopeptidase activity, malate metabolic process, coenzyme A biosynthetic process), transcription/RNA processing/translation (e.g., rRNA binding, piRNA processing, mRNA binding, regulation of gene expression, U3 snoRNA binding, RNA helicase activity, amino acid binding, regulation of DNA-templated transcription, RSC-type complex), and various other processes (e.g., 3-hydroxyacyl-CoA dehydratase activity, fatty acid elongation, alpha-catenin binding, lipid homeostasis, phosphatidic acid transfer activity, interleukin-17 receptor activity) (Fig. S14c; correlation value 0.48).

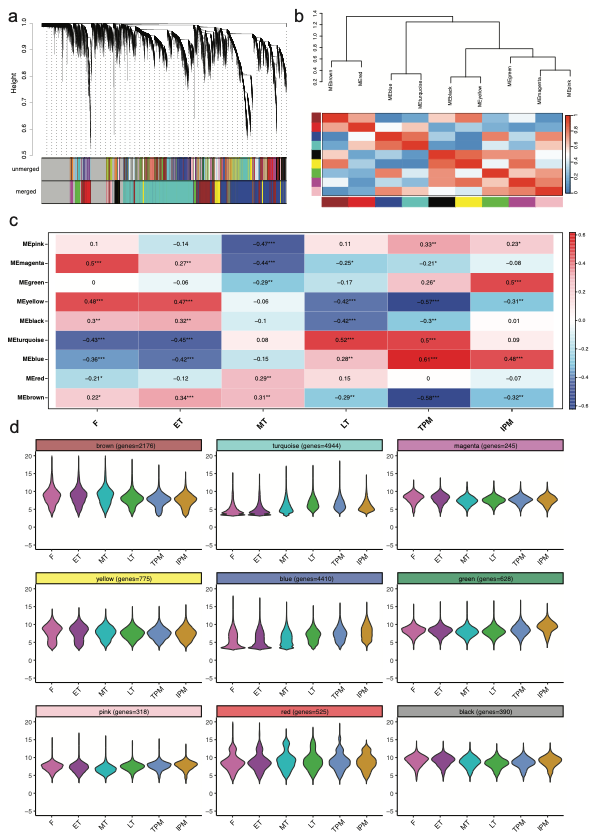

**Figure S14.** **WGCNA module generation and gene mapping results**. a) Cluster dendrogram showing unmerged (=69 modules + unassigned genes in grey module) and merged modules (=9 modules + unassigned genes in grey module); b) module-module correlation dendrogram and heatmap; c) module-trait correlation heatmap, with most significant correlations shown with three asterisks; d) Violin plots of expression across traits for genes assigned to each module (variance stabilised DESeq2 counts).

r
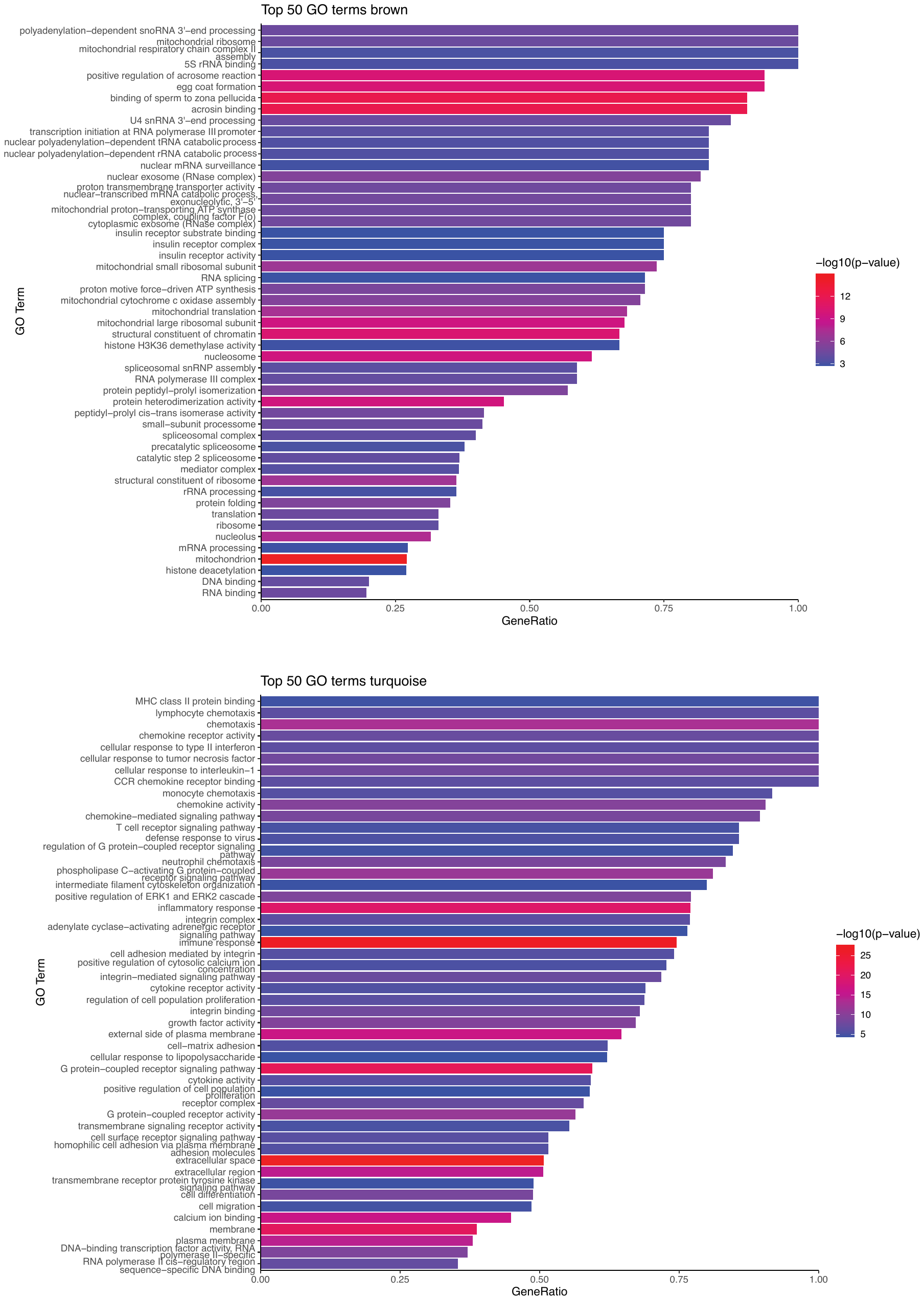

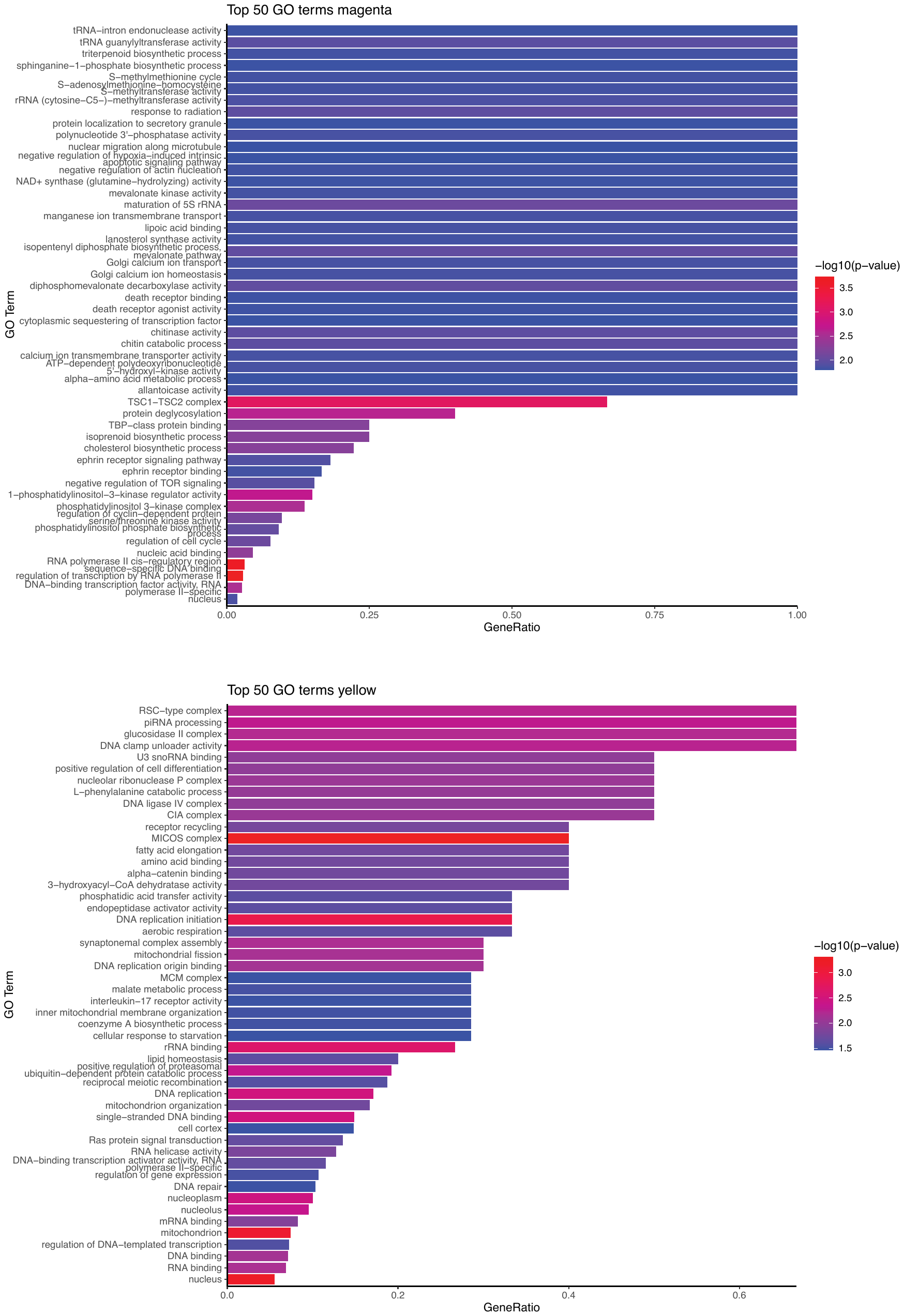

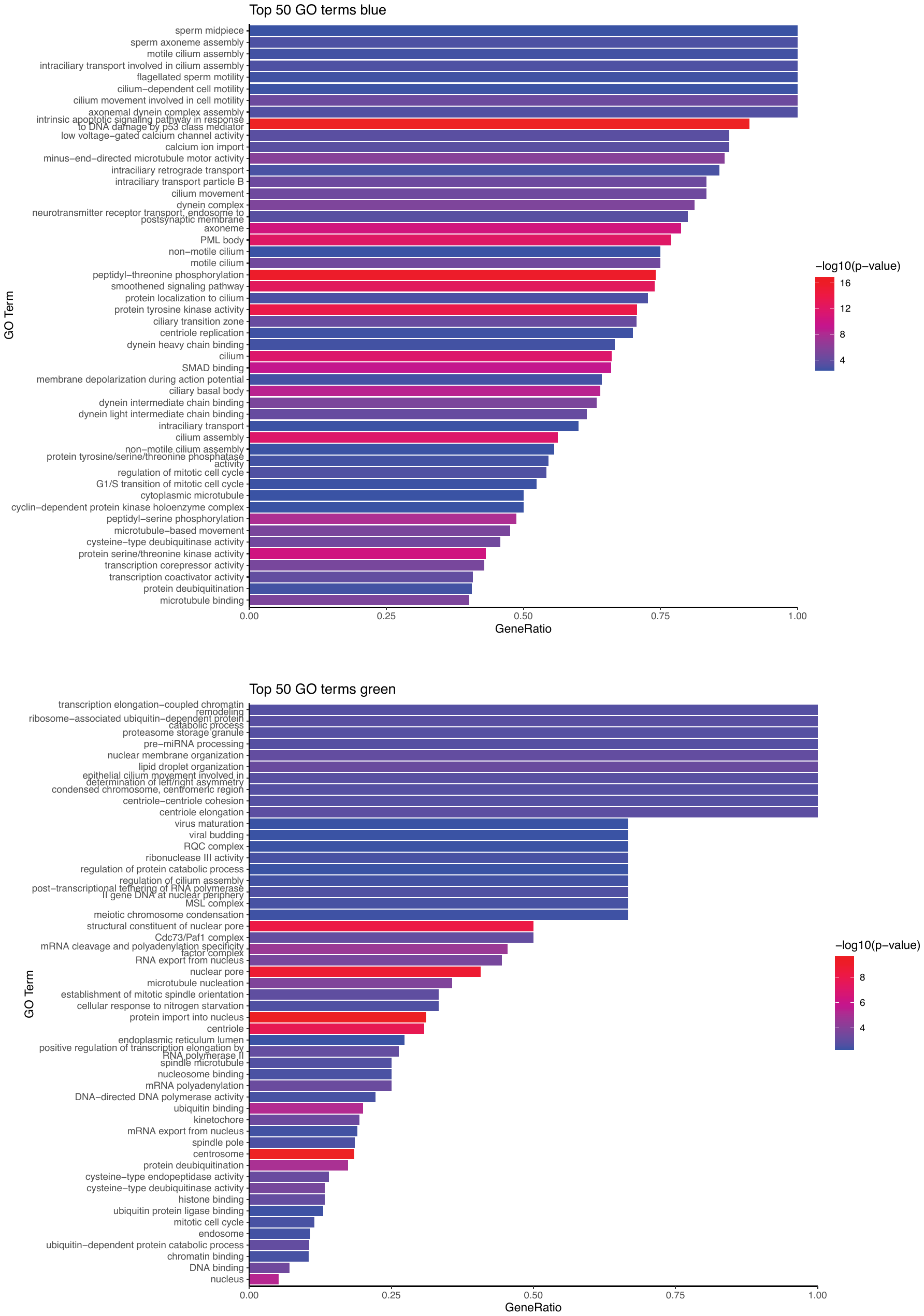

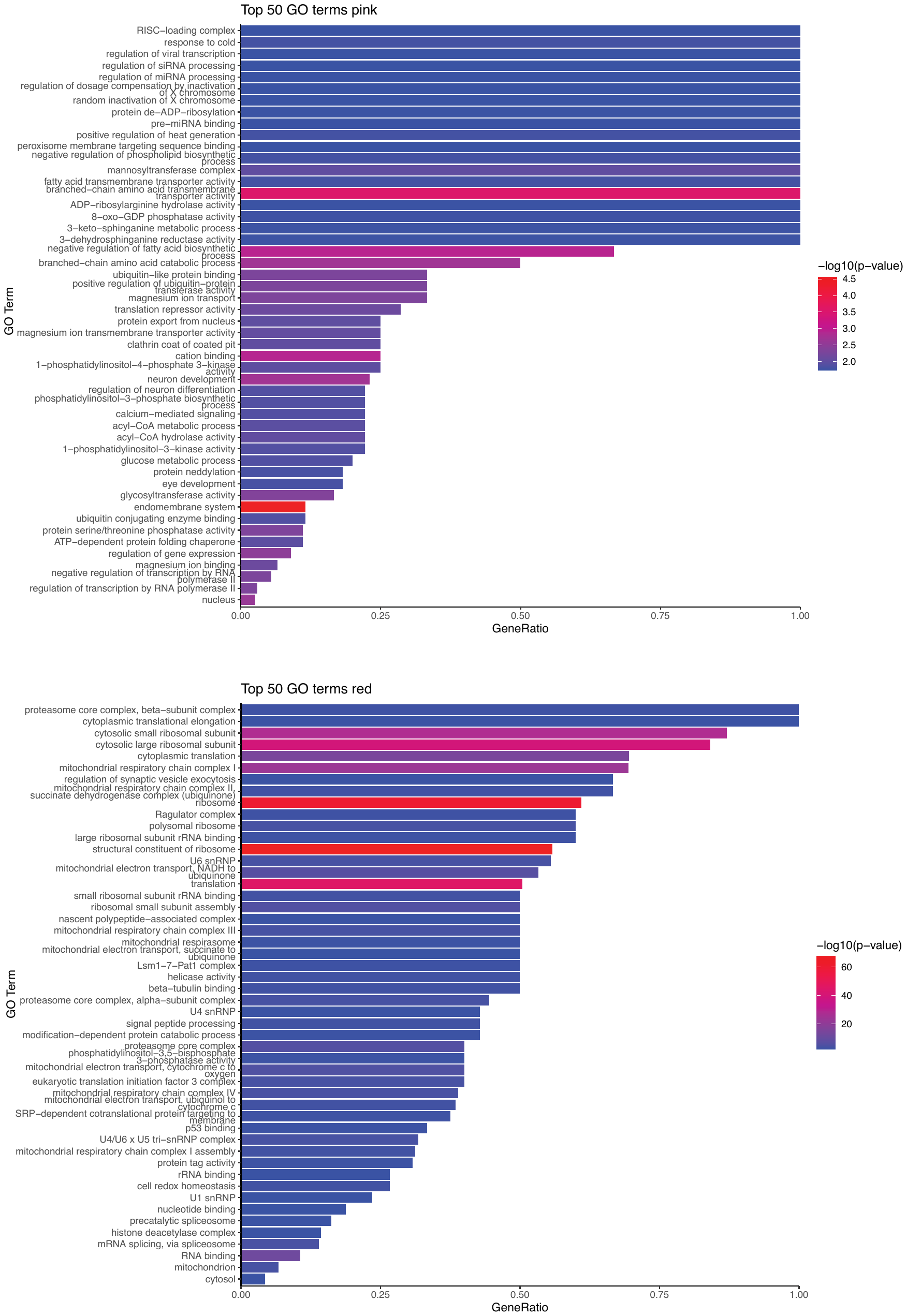

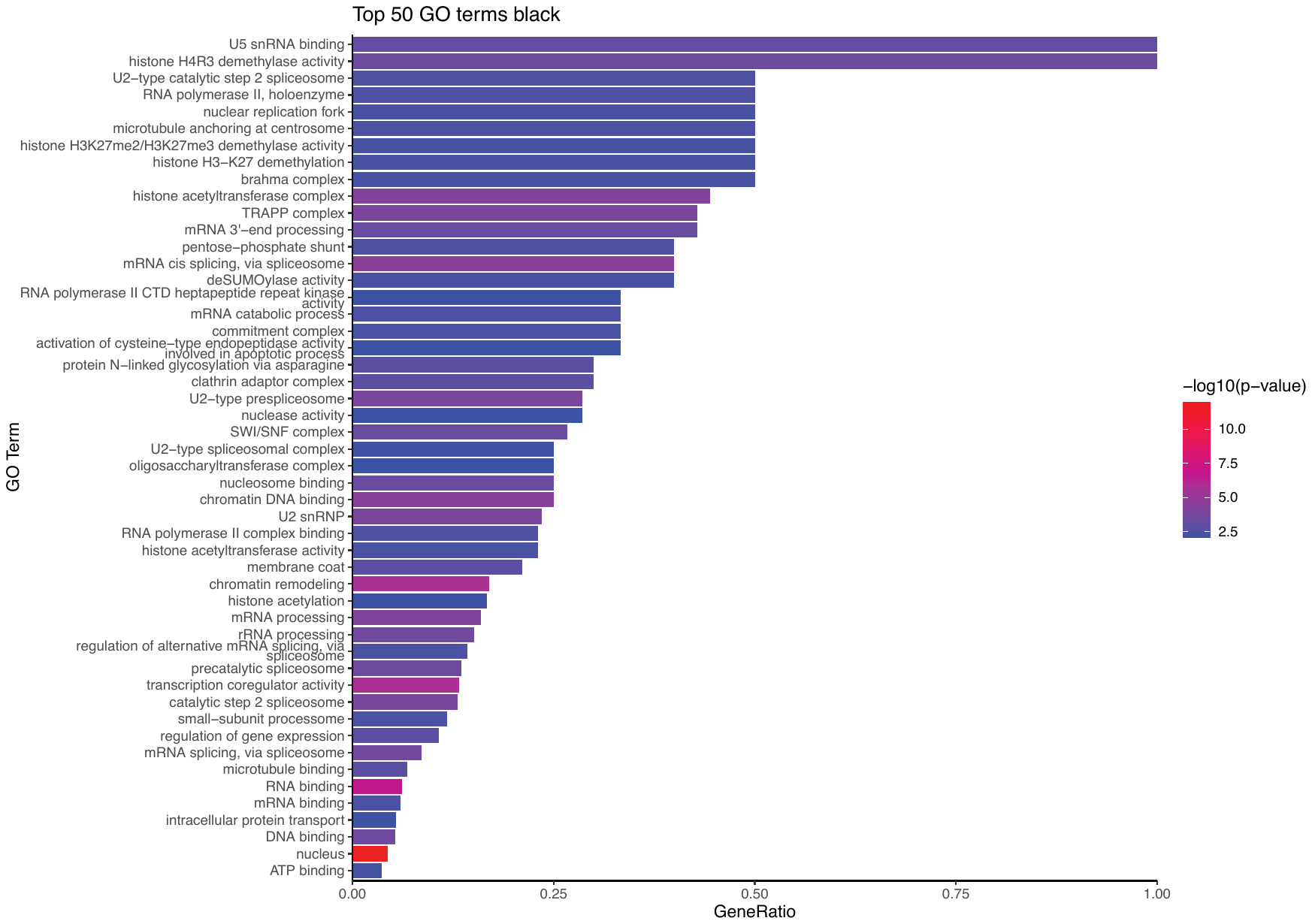
**Figure S15**. Top 50 enriched GO terms in genes designated to each module, in arbitrary order: brown, turquoise, magenta, yellow, blue, green, pink, red, black. Gene ratio calculated from numDEInCat/numInCat.

#### 6b. WGCNA: gene-trait correlation

During WGCNA, all genes are assigned a correlation value (and P-value) associated with each trait. These correlation values were then used to sort and select the top 30 most positively and negatively correlated genes with each trait. Some traits were very highly correlated with some genes, others less so. These are explored below.

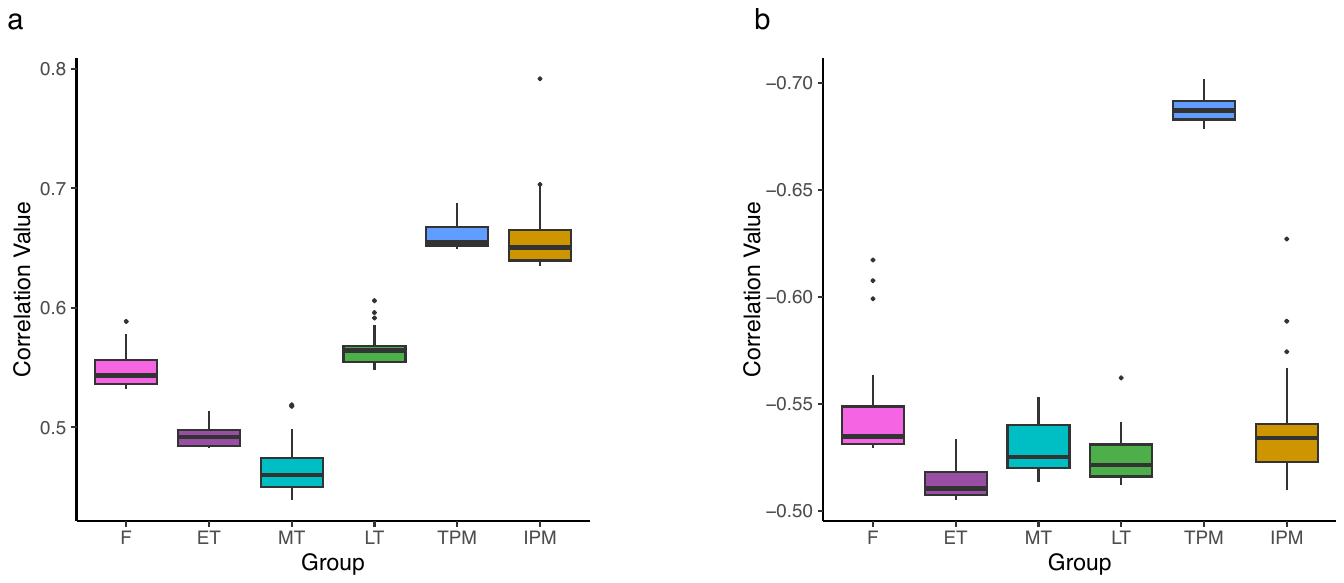

**Figure S16**. Distribution of correlation values for the top 30 genes a) positively and b) negatively correlated with each of the traits. Masculinised traits show the strongest correlations for the top 30 genes, in particular for genes that are negatively correlated with TPM. In general, the highest correlations of any genes with any one trait are not particularly high, with the highest single correlation being one gene (LOC117825029 - uncharacterized) seen as highly positively (0.79) correlated with IP males. Regardless, all correlations have p-values <0.001 (see Supplementary Table S11 for gene list, with corresponding correlation values used here and p-values. These genes are highlighted in blue in the table).
